# Targeted Tumor Microenvironment Delivery of Floxuridine Prodrug *via* Soluble Silica Nanoparticles in Malignant Melanoma as a Model for Aggressive Cancer Treatment

**DOI:** 10.1101/2025.03.14.643079

**Authors:** Andrés Ramos-Valle, Arnau Domínguez, Natalia Navarro, Ana Márquez-López, Anna Aviñó, Ramon Eritja, Carme Fàbrega, Lorena García-Hevia, Mónica. L. Fanarraga

**Affiliations:** The Nanomedicine Group, Institute Valdecilla-IDIVAL, 39011 Santander, Spain; Molecular Biology Department, Faculty of Medicine, Universidad de Cantabria, 39011 Santander, Spain; Dpt. Surfactants & Nanobiotechnology, Institute for Advanced Chemistry of Catalonia (IQAC), CSIC, 08034-Barcelona, Spain; CIBER-BBN Networking Centre on Bioengineering, Biomaterials and Nanomedicine, 08034-Barcelona, Spain

**Keywords:** malignant melanoma, VEGF receptor, TEM8, therapeutic oligonucleotide, silica nanoparticles, targeted delivery, tumor neovasculature

## Abstract

Malignant melanoma presents a significant challenge in oncology due to its aggressive nature and high metastatic potential. Conventional systemic treatments often fail to effectively reach tumor sites, limiting their therapeutic impact. This study introduces a groundbreaking triple-strategy approach for treating malignant melanoma. We developed a novel prodrug, an oligonucleotide, comprising 10 units of Floxuridine (5-fluoro-2’-deoxyuridine) (FdU) nucleoside antimetabolites, to enhance half-life and reduce rapid metabolism. Encapsulated in soluble colloidal silica nanoparticles, this compound is protected and directed towards tumor neovasculature precursor endothelial cell receptors, ensuring local delivery. The strategy focuses on releasing the prodrug in the tumor microenvironment, aiming to eradicate both melanoma cells and their supportive structures. Efficacy was demonstrated in cell culture studies and preclinical models of malignant melanoma, showing a remarkable 50% reduction in tumor size after just three intravenous treatments. These findings underscore the transformative potential of targeting endothelial cell membrane proteins for drug delivery. Our study paves the way for innovative targeted therapies, promising significant advancements in treatment strategies and improved outcomes for patients with metastatic cancers.

**Key Points:**

1. Triple-strategy for treating melanoma: FdU_10_ prodrug, silica nanoparticle and targeted delivery.
2. Oligonucleotide prodrug (Floxuridine units) enhances half-life and reduces metabolism.
3. Soluble silica nanoparticles protect therapeutic FdU_10_ from nucleases and decorated with protein ligands are directed to tumor neovasculature endothelial cells.
4. Significant 50% tumor reduction in preclinical melanoma models after systemic administration with targeted therapies.

## 1. Introduction

Resistance to chemotherapy poses a significant challenge for the systemic treatment of cancer. Despite advancements, many patients experience reduced drug efficacy as cancer cells develop drug resistance. This often necessitates increasing the drug dosages to levels that can threaten a patient’s life. This underscores the urgent need for strategies to overcome resistance and enhance therapeutic outcomes.

5-Fluorouracil (5-FU) is an essential component of cancer chemotherapy. This drug is effective against a variety of malignancies by inhibiting DNA and RNA synthesis in rapidly dividing cancer cells.^[1]^ However, like many other drugs, the short half-life and rapid metabolism of 5-FU require frequent dosing to maintain therapeutic levels and close monitoring of liver and kidney function to minimize side effects. ^[2,3]^

Various strategies have been explored to enhance the therapeutic outcomes and counter-resistance mechanisms of 5-FU. The combination of this drug with folic acid, leucovorin, irinotecan, and oxaliplatin,^[4]^ or the encapsulation of 5-FU in carriers such as alginate, chitosan, and carbon nanotubes has shown promising potential.^[5–10]^ While these approaches are hopeful, they present some limitations including drug-loading constraints and the risk of premature degradation. A recent advance involves the synthesis of oligonucleotide strands containing multiple units of 5-fluoro-2’-deoxyuridine (FdU), demonstrating superior efficacy compared to 5-FU alone.^[11,12]^ Notably, the oligonucleotide, composed of 10 units of Floxuridine (FdU_10_), has shown improved *in vitro* efficacy when incorporated into DNA-based nanoscaffolds.^[12–14]^ Here we explored the potential of this remarkable compound to enhance therapeutic efficacy through refined encapsulation and targeting strategies, while also addressing potential side effects. To counteract rapid elimination, and shield the drug from degradation, we encapsulated the compound within a novel type of soluble colloidal silica nanoparticles which have been recently validated for the delivery of DNA.^[15,16]^ This approach not only establishes a robust platform for drug delivery but also enables customized ligand-protein attachment directly to the nanoparticles, thereby improving delivery to tumoral tissues.

To achieve this, we have developed two novel protein ligands tailored to bind receptors on tumor vascular endothelial cells. This approach circumvents targeting cancer cells, which can develop resistance due to genetic instability and mutational changes that make them unrecognizable to engineered ligands. In contrast, endothelial cells in the tumor microenvironment (TME) are more accessible and play a pivotal role in tumor growth. Concentrating nanomedicines in these cells aids in releasing the prodrug upon dissolution of the nanoparticles in the surrounding tumor environment. This targeted release mechanism is expected to effectively target and eliminate both melanoma cells and their supportive structures.

## 2. Results

### 2.1. Synthesis and characterization of the FdU_10_Cy5 oligonucleotide

The synthesis of the FdU_10_Cy5 oligonucleotide process involved the preparation of the oligonucleotides using a solid-phase assembly of several units of FdU phosphoramidites obtaining the FdU_10_ oligomer, as described in the experimental section (**Figure 1A**). Oligonucleotides were cleaved from the solid support using ammonia treatment. The purity of the oligonucleotides was assessed using High-Performance Liquid Chromatography (HPLC), which confirmed the presence of a predominant product. As an inactive negative control for the study, we synthesized an oligonucleotide consisting of 10 thymidine residues (T_10_). Both oligonucleotides were synthesized *in-house* following standard phosphoramidite solid-phase chemistry as previously described and characterized.^[17]^ The identity of the oligonucleotides was confirmed by MALDI-TOF analysis (Table 1, Figure S1A and Figure S1B). To enable tracking of the oligonucleotides *in vitro* and *in vivo*, FdU_10_ and T_10_ oligonucleotides were labeled with Cyanine 5 dye (Cy5) at the 3’ end.

**Figure 1.**
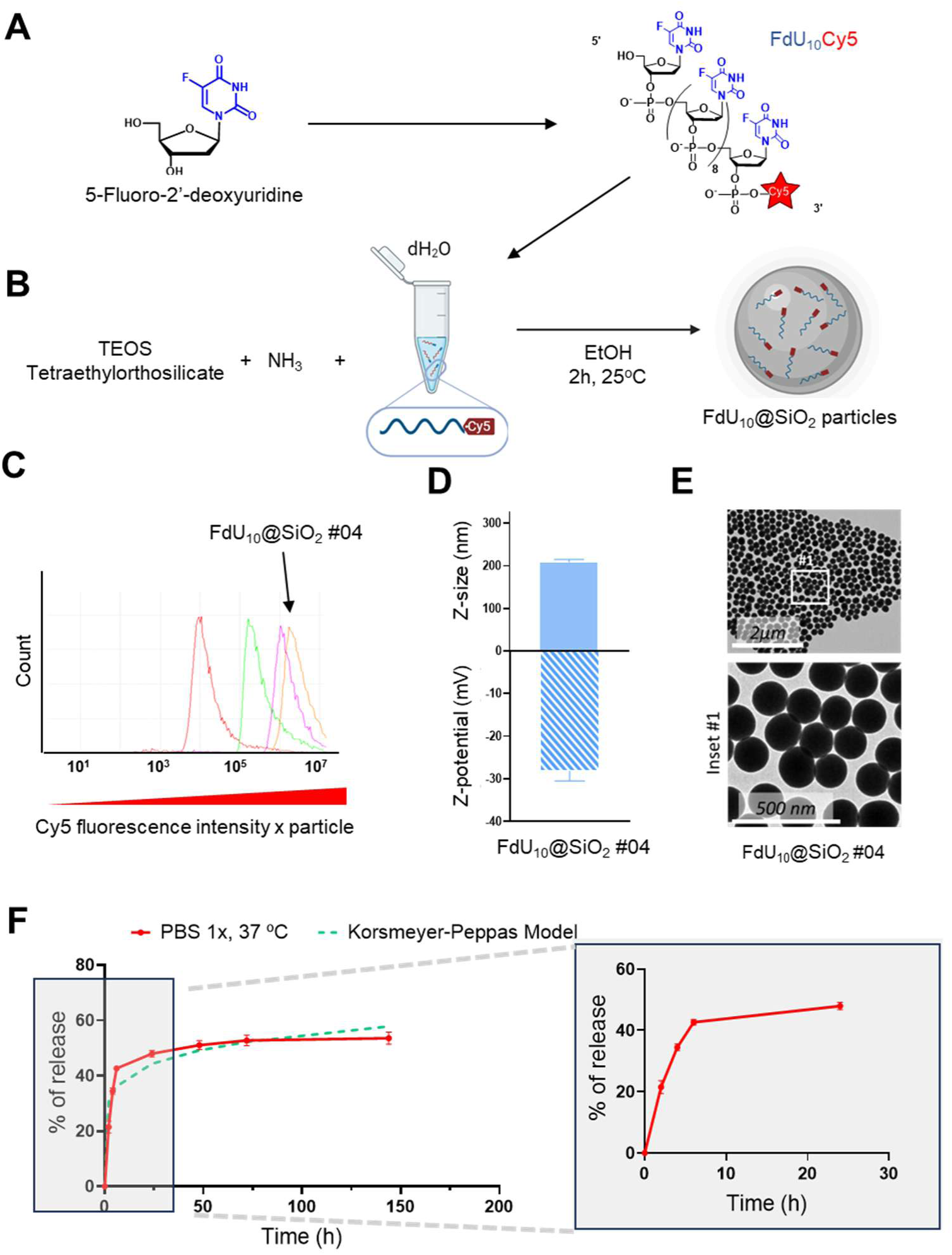
Nanoparticle synthesis and characterization. **A.** Schematic representation of the chemical synthesis of the FdU_10_ oligonucleotide labeled with Cy5. **B.** General diagram of the synthesis of FdU_10_@SiO_2_ particles using a modified Stöber method. **C.** Overlaid histograms of Cy5 intensity per each nanoparticle synthesis analyzed by flow cytometry (10,000 events per sample). FdU_10_@SiO_2_ #04 nanoparticles are indicated (orange). **D.** Z-size (nm) and ζ-potential (mV) of FdU_10_@SiO_2_ #04 particles. **E**. TEM images of FdU_10_@SiO_2_#04 particle morphology. Inset #1 showing an augmented caption of the particle morphology and the scale adjustment based on the raw image. **F.** Profile of the release of the prodrug from FdU_10_@SiO_2_ #04 particles in PBS at 37°C for 6 days (in red). The dashed green line represents the Korsmeyer-Peppas model fitting to the release kinetics.

**Table 1.**
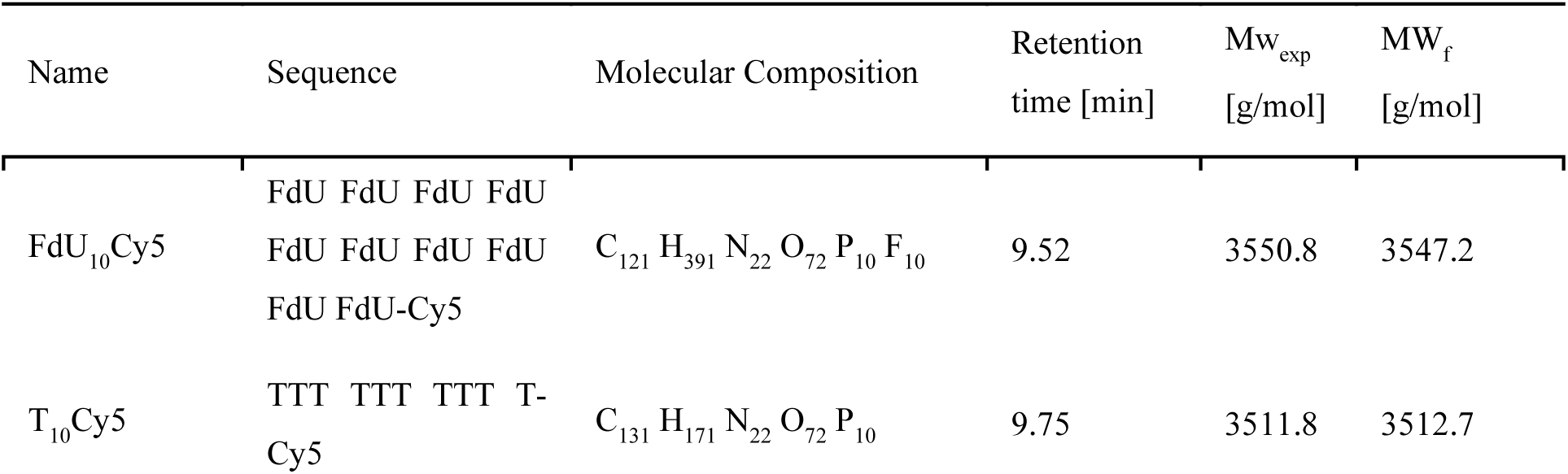
Sequences and Molecular Characterization of Synthesized FdU_10_-Cy5 and T_10_-Cy5 Oligonucleotides. The expected (exp) and found (f) molecular weights are provided for comparison, ensuring accuracy in the synthesis process.

### 2.2. FdU_10_Cy5 encapsulation and release from FdU_10_@SiO_2_ nanoparticles

To protect the oligonucleotide from elimination and degradation we opted to encapsulate the FdU_10_Cy5 prodrug within soluble colloidal silica nanoparticles (Figure 1B). As demonstrated in our previous investigations, this encapsulation system effectively safeguards DNA integrity and liberates nucleic acids upon dissolution of the silica coating under physiological conditions, both *in vitro* and *in vivo*.^[15,16]^

The FdU_10_Cy5 silica nanoparticles (hereafter FdU_10_@SiO_2_) were prepared following a previously established protocol based on a modified Stöber procedure.^[15,16]^ The single-stranded nature and small size (10 residues) of the FdU_10_ oligonucleotides necessitated fine-tuning of the formulation. Therefore, we tested different stoichiometric ratios (Table S1). Among the various formulations evaluated, we observed that increasing the ethanol content promoted the synthesis of smaller and more uniformly sized nanoparticles (Figure S2). This adjustment also resulted in higher encapsulation efficiency of FdU_10_Cy5, as demonstrated by flow cytometry analysis of the loaded nanoparticles (Figure 1C and Figure S3). Based on these results, the formulation FdU_10_@SiO_2_#04 (henceforth referred to as FdU_10_@SiO_2_) was selected for the trials. These optimized nanoparticles had a diameter of 190 nm by TEM, a ζ potential of around −27 mV, and a PDI of 0.08 ± 0.06 (Figure 1D and Figure 1E). Similar results were observed for the control nanoparticles loaded with the T_10_-Cy5 oligonucleotide (referred to as T_10_@SiO_2_) (Figure S4).

Direct measurement of the encapsulated oligonucleotide within the FdU_10_@SiO_2_ silica nanobeads was conducted using thermogravimetric analysis (TGA). The results indicated that 97.7% of the FdU_10_Cy5 total mass added to the synthesis was entrapped in the nanoparticles (Figure S5). This signifies that the embedding efficiency of the FdU_10_@SiO_2_ nanoparticles was 97.1% of the initial oligonucleotide included in the mixture, with a 0.44% loading capacity explained by the compact morphology of the particles and the higher density of silica.

To complement this study, we assessed the efficiency of oligonucleotide release upon the dissolution of the FdU_10_@SiO_2_ nanoparticles *in vitro* upon incubation in phosphate-buffered saline (PBS) at 37°C.^[15,16]^ Using fluorimetry, we quantified the release kinetics of the FdU_10_Cy5 oligonucleotide into the media (Figure S6). Figure 1F illustrates that approximately 40% of the embedded oligonucleotide was released within the initial 6 hours. Following this, a sustained and prolonged pattern of drug release was observed, persisting for over 6 days. This data was analyzed using the Korsmeyer-Peppas model, revealing a consistent fit with this model, characterized by a release exponent value of 0.14 and an R^2^ value of 0.94. The drug release mechanism was determined to be diffusion (Fickian model), with a slope of <0.5 (Table S2).

### 2.3. Comparative evaluation of free and encapsulated FdU_10_Cy5 in malignant melanoma cell cultures

Malignant melanoma is a major challenge in oncology due to its resistance and aggressiveness. With drug resistance mechanisms involving complex pathways, combating melanoma becomes increasingly difficult as it progresses to the metastatic stage. The grim reality is reflected in the statistic that 30% of patients succumb to the disease within five years.^[18,19]^ This underscores the urgent need for continued research and innovative treatment approaches to improve patient outcomes.^[20–23]^

To assess the impact of the FdU_10_ oligonucleotide prodrug on B16-F10 malignant melanoma cells, we undertook a comparative analysis concentrating on the efficiency of the decapsulated FdU_10_Cy5 oligonucleotides. We specifically evaluated the efficacy and effects of FdU_10_@SiO_2_ nanoparticles compared to the free drug, alongside a control group treated with the inactive control T_10_Cy5, both in free form and encapsulated.

Flow cytometry analysis and fluorescence microscopy imaging confirmed the internalization of the nanoparticles by the cells within 2 h of treatment (Figure 2A and Figure 2B, arrow). Flow cytometry analysis quantitatively demonstrated that the intracellular Mean Fluorescent Intensity (MFI) of the Cy5 fluorophore, representing the total uptake of encapsulated FdU_10_-Cy5 in FdU_10_@SiO2 nanoparticles, was significantly higher compared to the free FdU_10_-Cy5. Specifically, the MFI was 16.6 times higher 2 hours after nanoparticle exposure and 38.5 times higher after 24 hours, highlighting the enhanced cellular uptake and retention of the nanoparticle-encapsulated drug over time. This trend was observed at all the time points analyzed (48 h and 72 h, Figure S7). Previous studies with similar nanoparticles have consistently demonstrated rapid cellular interactions, with internalization occurring within a few hours.^[24,25]^ The intracellular localization of the compounds was confirmed using single-plane confocal microscopy imaging. Melanoma cells treated with free FdU_10_Cy5 oligonucleotide showed Cy5 fluorescence at the perinuclear cytoplasmic region, confirming oligonucleotide internalization, whereas FdU_10_@SiO_2_ nanoparticles were scattered throughout the cytoplasm of the cell (Figure 2B and Figure 2D, arrows). At this time point, no detectable fluorescent drug release was observed using confocal microscopy.

**Figure 2.**
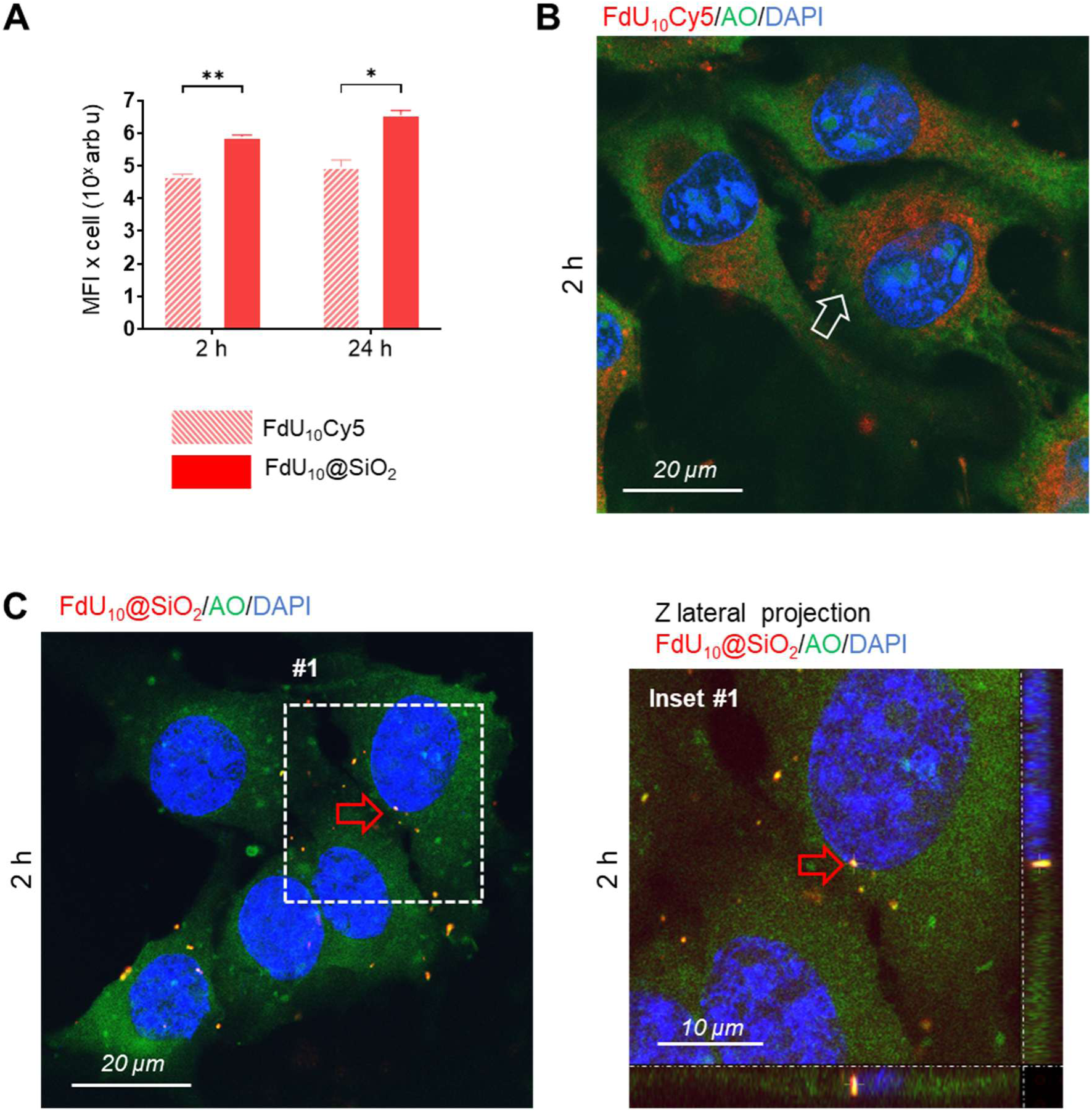
Cellular capture of the free/encapsulated prodrug. **A.** Mean fluorescence intensity (MFI) of Cy5 in melanoma cells treated with free FdU_10_-Cy5, and FdU_10_@SiO_2_ particles at 2 h and 24 h (Data are presented as mean values ± S.D, *n*=3 (10,000 cells per replica) * *p<*0.05, \*\**p<*0.01, *t*-test comparisons, see Figure S7). Figures B and C show single Z-plane confocal microscopy images of melanoma cells 2 hours after the indicated treatments. **B**. The free FdU_10_-Cy5 oligonucleotide (red) is distributed primarily in the perinuclear cytoplasmic region (white arrow). **C**. FdU_10_@SiO_2_ particles display delayed intracellular drug release. Inset #1 shows lateral projections of an intracellular nanoparticle (red arrow), with several other nanoparticles (orange) visible in the same Z-plane. Cy5 fluorescence is depicted in red, while the cell cytoplasm and nucleus are stained green and blue, respectively.

At 72 h, the cells treated with both, FdU_10_@SiO_2_ or free FdU_10_Cy5 oligonucleotide exhibited noticeable effects, including increased nuclear and cytoplasmic sizes compared to the untreated cells and T_10_@SiO_2_ nanoparticle-treated controls (Figure 3A and Figure 3B). At this time point, the accumulation of Cy5 fluorescence in the cytoplasm confirmed the presence of a substantial amount of free and encapsulated intracellular oligonucleotides (Figure 3A, arrows).

**Figure 3.**
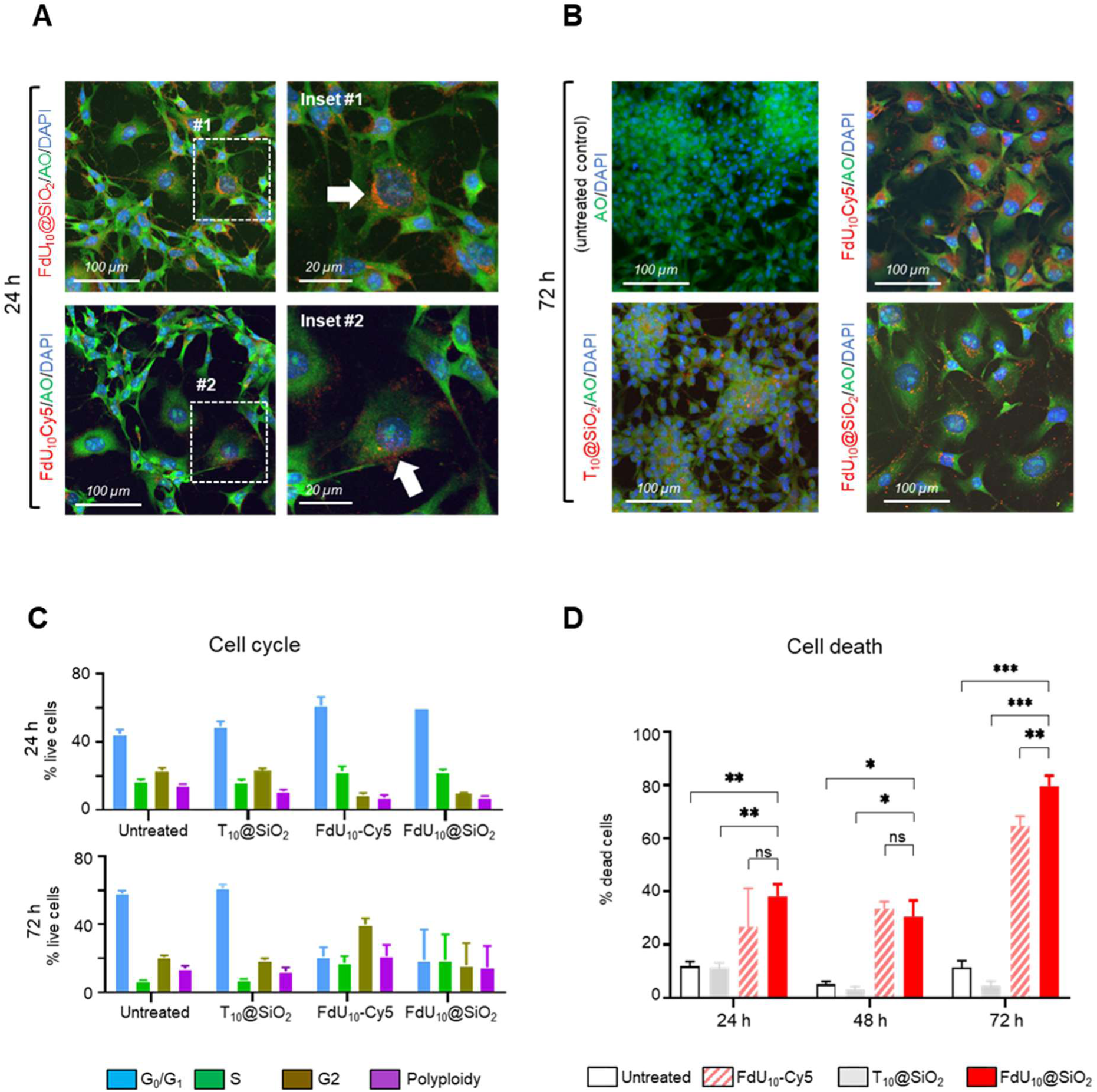
*In cellulo* effects of FdU_10_@SiO_2_ particles and controls. **A-B.** Confocal microscopy images of melanoma cells at 24 and 72 hours post-treatment. Cells treated with PBS and T_10_@SiO_2_ nanoparticles exhibited significant proliferation. In contrast, a notable increase in cell size was observed in cultures treated with either the free FdU_10_ or the silica-encapsulated FdU_10_ prodrug. **C**. Flow cytometry analysis of the cell cycle in melanoma cultures at 24 and 72 hours post-treatment revealed significant abnormalities following treatment with the drug alone or FdU_10_@SiO_2_ nanoparticles. **D**. Flow cytometry quantification of cell death in untreated (control) melanoma cells versus those exposed to T_10_@SiO_2_, FdU_10_Cy5, and FdU_10_@SiO_2_. Data are presented as mean values ± S.D. (*n*=3), with \**p*<0.05, \*\**p*<0.01, and \*\*\**p*<0.001 indicating statistical significance. Statistical analysis was performed using one-way ANOVA with multiple *t*-test comparisons, with a total of 10,000 cells analyzed per flow cytometry replicate.

Flow cytometry analysis revealed significant cell cycle disruptions in cells treated with both free and encapsulated FdU_10_-Cy5, showing a clear G0/G1 phase arrest at 24 hours (Figure 3C and Figure S8). This blockage aligns with the inhibition of DNA synthesis, preventing the cells from entering the S phase. By 72 hours, the cell cycle showed further dysregulation, with a decrease in G0/G1 phase cells and an uncoordinated distribution across other phases. This pattern is consistent with the mechanism of action of 5-FU,^[1]^ suggesting that the surviving cells transition into a non-proliferative state, as corroborated by Ki67 and p21 immunostaining results (Figure S9).

Finally, to demonstrate the cytotoxic effect of these formulations, we quantified tumor cell death (Figure 3D). At 24 hours, cells treated with free FdU_10_Cy5 oligonucleotide or FdU_10_@SiO_2_ nanoparticles showed death rates of 23% and 33%, respectively compared to untreated control cultures. After 72 hours, cell mortality increased to 65% and 79%, respectively. In contrast, cells exposed to the T_10_@SiO_2_ control nanoparticles showed no significant morphological or cell cycle changes. This demonstrates that the observed effect is due to the FdU_10_Cy5 oligonucleotide and not to the nanoparticles themselves. Together, these findings demonstrate that, at least in cell culture, the effect of the FdU_10_Cy5 prodrug can be significantly enhanced when encapsulated in silica nanoparticles.

### 2.4. Enhanced antitumor efficacy of locally delivered FdU_10_@SiO_2_ nanoparticles

After the validation of the FdU_10_@SiO_2_ nanoparticles *in cellulo*, we proceeded to investigate the *in vivo* effects of the treatments through local intratumoral injection. This experiment aimed to evaluate the antitumor efficacy of both free and encapsulated FdU_10_ within the tumor microenvironment, independent of their targeting efficiency. The objective was to demonstrate the effective release of the prodrug from the silica particles, the subsequent activation of the drug within the tumor, and its antitumoral effects, regardless of any targeting considerations. To this end, we used immunocompetent tumor-cell-transplanted mice as a preclinical model of malignant melanoma, (Figure 4A), a model that has been widely validated in the literature.^[10,26–28]^ Transplanted cancer cells produce solid-pigmented melanoma tumors that are easy to monitor, isolate, and assess through anatomopathological examination, ensuring good reproducibility. These *in vivo* studies enable exploration of the complex interplay between the TME, tumor cells, and tumor-targeted nanomedicines, providing insights into cancer progression and treatment efficacy.

**Figure 4.**
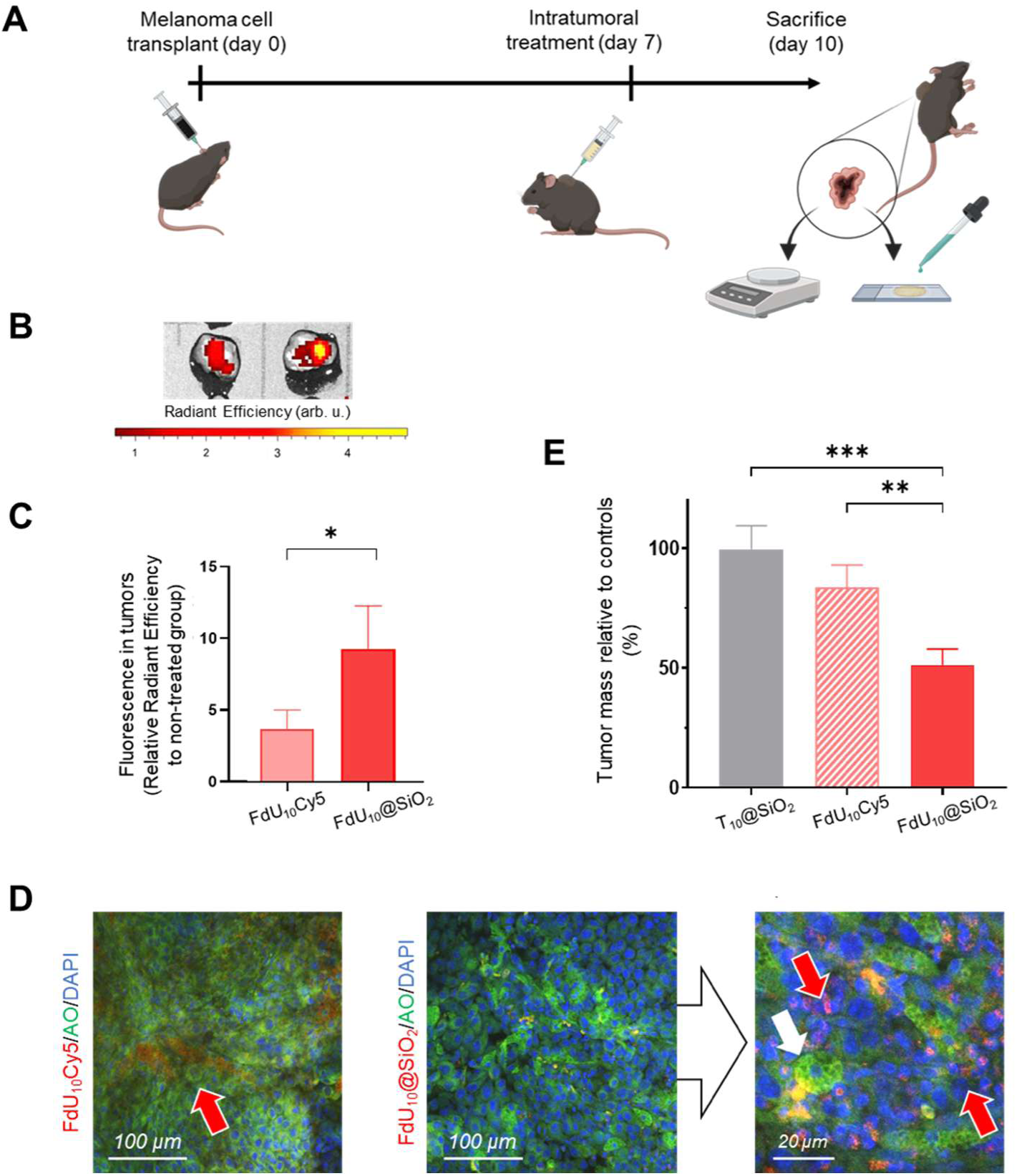
*In vivo* intratumoral efficacy of the encapsulated compound. **A.** General scheme and timeline of the melanoma model developed to evaluate the intratumoral efficacy of the treatments. **B**. *In toto* Cy5 fluorescence imaging in whole tumors. **C**. Quantification of Cy5 fluorescence detected in the tumors at sacrifice captured with an IVIS® imaging system (relative radiant efficiency for each *n=7* per group, \**p<*0.05, *t*-test analysis). **D**. Confocal microscopy images of the tumor tissues exposed to the indicated treatments. The fluorescence of the Cy5-labeled prodrug is shown in the red channel (red arrows). Cell cytoplasms and nuclei are shown in green and blue, respectively. A white arrow indicates a macrophage within the tumor that does not contain detectable nanoparticles. **E**. Quantification of relative tumor weights after intratumoral treatments compared to the PBS-treated control group (Data are presented as mean values ± S.D, *n=*10 mice/group, \**p<*0.05, \*\**p<*0.01, \*\*\**p<*0.001, One-way ANOVA with multiple *t*-test comparisons).

The treatments comprised a single injection of either the free oligonucleotide FdU_10_Cy5, the encapsulated form in FdU_10_@SiO_2_ nanoparticles, or the negative controls (PBS or T_10_@SiO_2_ nanoparticles). All the treatments were administered at identical total oligonucleotide doses. Mice were sacrificed three days post-injection (day 10 post-transplant).

The local availability of the prodrug in the tumors was monitored for 72 h after treatment using an IVIS® biofluorescence imaging system (Figure S10). Quantification of total Cy5 fluorescence at the tumoral tissues revealed a significant 3-fold increase in FdU_10_Cy5 when encapsulated in nanoparticles compared to the free oligonucleotide (Figure 4A and Figure 4C). Confocal microscopy of the tumor tissue showed that the nanoparticles were visible throughout the tumor microenvironment compared to the free prodrug, which was diffusely distributed in the tissue (Figure 4D, arrows). This suggests improved local drug retention. Consistent with this, tumors treated with FdU_10_Cy5 decreased in size by almost 20% compared to PBS-treated controls, whereas those treated with FdU_10_@SiO_2_ were reduced by more than half (*p*=0.009, *n=*15) (Figure 4E).

As in the *in cellulo* study, these results demonstrate that the efficacy of the FdU_10_Cy5 oligonucleotide is significantly improved when administered in the encapsulated form. The high vascularity of the tumor likely contributes to the systemic dispersion and degradation of the free oligonucleotides. In contrast, the FdU_10_@SiO_2_ nanoparticles enabled precise deposition and gradual local prodrug release, culminating in a remarkable *ca*. 50% reduction in the tumor mass following a single injection.

### 2.5. Systemic FdU_10_@SiO_2_ nanoparticle targeting to the neovasculature results in significant antitumoral effects

Targeting nanoencapsulated chemotherapeutic drugs to the TME is pivotal in oncology. Unlike traditional methods focusing on cancer cell receptors prone to resistance, TME cells offer stable targets accessible for nanoparticle delivery. Here, our approach focuses on targeting neovascular cells using FdU_10_@SiO_2_ nanoparticles. The hypothesis driving this strategy posits that delivering nanoparticles to surface receptors of endothelial cells enables their dissolution and gradual drug release in the vicinity. This process aims to inhibit blood vessel formation and induce cell death in the surrounding tumor microenvironment, indirectly targeting cancer cells. Consequently, this approach exerts a cytotoxic effect that affects both melanoma cells and supportive TME.^[29,30]^

To validate the hypothesis, we explored two distinct tumor neovasculature-targeted ligands. One approach involved targeting the vascular endothelial growth factor receptor (VEGFR), based on its documented success in previous studies for achieving significant antitumoral effects with nanomaterials.^[31–36]^ As a targeting agent we have selected the peptide ligand ATWLPPR, which is an effective antagonist of VEGFR.^[37]^ For functionalization purposes, we genetically fused the peptide to the carboxyl terminus of GFP, resulting in a protein termed GFP:VEGFRbp (Figure 5A and Figure S11A).

**Figure 5.**
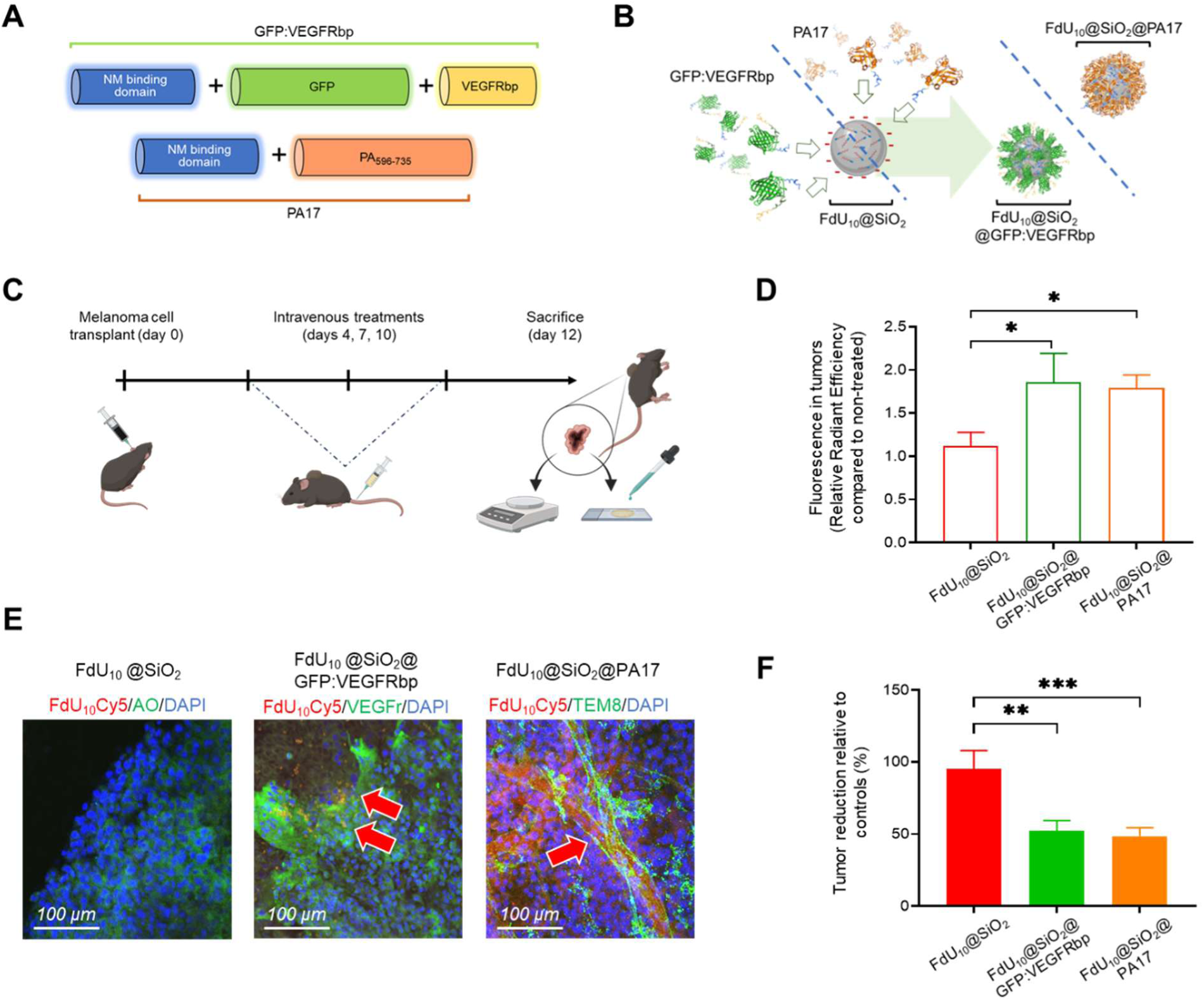
Neovasculature-targeted treatments *via* intravenous administration. **A.** Schematic representation of the protein design for GFP:VEGFRbp and PA17. **B**. Diagram of the functionalization process to produce FdU_10_@SiO_2_@GFP:VEGFRbp and FdU_10_@SiO_2_@PA17 nanoparticles **C**. General schematic and timeline for tumor generation and intravenous administration of the experimental treatments to melanoma-bearing mice. **D**. Quantification of Cy5 fluorescence detected in the tumors at sacrifice (day 12), captured with an IVIS® biofluorescence imaging system, with relative radiant efficiency for each treatment group. Data are presented as mean values ± S.D. (*n*=5), with significance indicated as \**p*<0.05 and \*\**p*<0.01, determined by One-way ANOVA with multiple *t*-test comparisons. **E**. Confocal microscopy proyection images of melanoma tumors treated with FdU_10_@SiO_2_, FdU_10_@SiO_2_@GFP:VEGFRbp, and FdU_10_@SiO_2_@PA17. VEGFR and TEM8 receptors are immunostained and visualized in the green channel, cell nuclei in the blue channel, and the prodrug in the red channel. **F**. Relative tumor masses of the groups treated with T_10_@SiO_2_, free FdU_10_-Cy5, and FdU_10_@SiO_2_ compared to the PBS-treated group (in %). Data are presented as mean values ± S.D. (*n*=10 mice per group), with significance indicated as \**p*<0.05, \*\**p*<0.01, and \*\*\**p*<0.001, analyzed using One-way ANOVA with multiple *t*-test comparisons.

In parallel, we engineered another neovasculature-targeted ligand inspired by the protective antigen protein (PA), which has a high affinity for the tumor endothelial marker 8 (TEM8) predominant in the tumor neovasculature. This receptor is abundantly expressed on the membranes of angiogenic endothelial cells, associated stromal cells, pericytes, cancer stem and invasive cancer cells, and immune cells such as macrophages and cancer-associated fibroblasts. It was also extraordinarily expressed in the murine malignant melanoma (Figure S12).^[38–40]^ As for VEGFR inhibitors,^[41,42]^ the blockade^[43]^ or knockout^[44–47]^ of TEM8 has been shown to effectively suppress tumor growth while sparing physiological angiogenesis. This approach presents a promising new direction in cancer therapy with minimal adverse effects.^[48–50]^ The engineered PA-inspired ligand, which we named PA17 (Figure 5A and Figure S11B), had a molecular weight of 17 kDa and had a high affinity for the TEM8 receptor.^[51]^

The two engineered ligand-proteins were produced recombinantly and purified according to standard procedures (Methods). They were subsequently used to electrostatically functionalize the FdU_10_@SiO_2_ and control T_10_@SiO_2_ nanoparticles following an established protocol (Figure 5B).^[52]^ The correct surface functionalization with the ligand proteins was confirmed using SDS-PAGE (Figure S13). The functionalized nanoparticles were named FdU_10_@SiO_2_@GFP:VEGFRbp and FdU_10_@SiO_2_@PA17.

The validation of these targeted nanomedicines was performed in preclinical immunocompetent animal models of malignant melanoma. Systemic treatment involved three intravenous injections on days 4, 7, and 10 post-transplantation, as detailed in Figure 5C (Methods). A total amount of 120 µg of the encapsulated FdU_10_Cy5 oligonucleotide was applied both, in bare nanoparticles (non-functionalized) as control, and particles functionalized with the targeting ligands. Regular monitoring and weighing of mice throughout the treatment period showed no evidence of any toxic effects (Figure S14). Treated mice were euthanized on day 12 post-transplantation.

A semi-quantitative assessment of oligonucleotide accumulation *in toto* using an IVIS® biofluorescence detection system confirmed a significant increase in FdU_10_Cy5 accumulation in tumors. This was the case for nanoparticles coated with both GFP:VEGFRbp and PA17 ligands, with 1.7 and 1.6 times more than naked FdU_10_@SiO_2_ particles, respectively (Figure 5D and S15). The effective delivery of the FdU_10_Cy5 fluorescent oligonucleotide was corroborated by confocal microscopy imaging of the fresh tumor stroma. Notably, distinct Cy5 fluorescence patterns indicating prodrug distribution were observed in tumors of mice treated with both targeted nanoparticle variants. In mice treated with FdU_10_@SiO_2_@VEGFRbp nanoparticles, prodrug accumulation appeared as red spots predominantly located near VEGFR-positive tissue. In contrast, tumors treated with FdU_10_@SiO_2_@PA17 nanoparticles showed increased fluorescence primarily within and around the periphery of the tumor vasculature (Figure 5E, arrows).

Quantification of the antitumoral effect revealed a significant 50% reduction in tumor size using targeted nanoparticles compared to PBS-treated controls (Figure 5F). The T_10_@SiO_2_ nanoparticles functionalized with the ligand proteins showed limited effectiveness in reducing tumor mass, despite some intrinsic ligand activity (Figure S16). Bare FdU_10_@SiO_2_ nanoparticles used as controls exhibited only a modest 5% reduction. These results highlight the significantly greater efficacy of the targeted encapsulated drug treatment.

Finally, to confirm the effectiveness of the targeting mechanism in reducing tumor neovasculature, we conducted histochemical evaluations of the tumoral tissues. As depicted in Figure S17, treatment with the two endothelial cell-targeting nanoparticles significantly reduced the intratumoral neovasculature in approximately a 75% reduction compared to control tumors treated with PBS or the non-functionalized FdU_10_@SiO_2_ nanoparticles.

## 3. Discussion

Systemic pharmacological treatments are effective but often have short durations of action and require high doses, leading to severe side effects. Nanoencapsulation of drugs offers a promising solution by enabling controlled release and prolonged drug performance. However, only 1-2% of administered nanomedicines reach the tumor,^[53,54]^ posing significant challenges in aggressive cancers like malignant melanoma.

A pivotal aspect of this study revolves around synthesizing and delivering antiproliferative oligonucleotides as a prodrug. Encapsulation within soluble silica nanoparticles protects the prodrug from degradation and swift elimination, while novel protein ligands of VEGF and TEM8 receptors precisely target these nanoparticles to the tumor neovasculature and TME. Upon reaching the tumor tissues, the FdU_10_ oligonucleotide undergoes decapsulation and exonuclease activation to release active 5’-phosphate nucleotide derivatives, thereby reducing potential systemic toxicity. This targeted approach aims to trigger an effect on both cancer cells and supportive microenvironmental cells.

The utilization of an immunocompetent (*wild type*) animal model has also been instrumental in enabling drug delivery strategies tailored to the cellular components of the TME. Most trials and proof-of-concept tests assessing the efficacy of nanomedicine have utilized nude animals with severely compromised immune systems. Hence, the effects observed for many drug nanodelivery studies in these animals are notably reduced when wild-type immunocompetent animals are employed.

In summary, the combination of FdU_10_ with silica particles and a targeted strategy directed at the tumor neovasculature has shown great potential. The achievement of 50% antitumor activity after only three intravenous treatments highlights the critical importance of targeting the microenvironment in aggressive cancer therapy. This study paves the way for more effective and less toxic treatment options for aggressive cancers such as malignant melanoma.

## 4. Conclusion

The developed treatment strategy for malignant melanoma has yielded remarkable outcomes through the integration of the potent FdU_10_ prodrug compound with a unique soluble silica nanoparticle encapsulation system and two distinct tumor endothelial cell-targeted ligands. This approach has led to a notable reduction in tumor size, approximately 50%, emphasizing the effectiveness of targeting cells within the TME rather than solely focusing on cancer cell receptors. These results establish a foundation for tailored therapies and bring hope to patients battling malignant melanoma and other aggressive metastatic cancers. By specifically targeting the neovasculature with FdU_10_-loaded nanoparticles, our study offers a promising avenue for future therapeutic interventions.

## 5. Methods

### Oligonucleotide synthesis and characterization

Briefly, oligonucleotide synthesis was performed at a 1 µmol scale using phosphoramidite solid-phase protocols as described previously.^[55]^ After ammonia deprotection, oligonucleotides were desalted using Sephadex G-25 columns and used without additional purification. The purity of the oligonucleotides was analyzed by HPLC and MALDI-TOF mass spectroscopy (more details in Supplementary Methods).

### Synthesis, characterization, and prodrug loading of FdU_10_@SiO_2_ nanoparticles

In a 1.5 ml microtube, the corresponding 4.5 µg of FdU_10_-Cy5 (see Table S2, for particles #4 = 4.5 µg) were dispersed in 18 µl of nuclease-free ddH_2_O (2.5 M). This FdU_10_-Cy5 solution was poured into a 1.5 ml microtube containing 236 µl of absolute EtOH (for particles FdU_10_@SiO_2_#4, rest see Table S1). The mixture was stirred at 1,200 r.p.m for 5 min. Subsequently, 17.3 µl of ammonia 25% (0.31 M) and 5.6 µl TEOS (0.06 M) were added to this mixture. The reaction mixture was then vortexed for 2 h at 1,200 r.p.m. FdU_10_@SiO_2_ spheres were centrifuged at 6,500 r.p.m, washed three times with absolute EtOH, and stored in 100 µl EtOH.

TEM images were obtained with a JEM1011 equipped with a high-resolution Gatan digital camera (JEOL) and analyzed using ImageJ software. DLS size distribution and Z-potential measurements were performed in triplicate using a Malvern Panalytical Ultra Zetasizer, with data presented as mean ± standard deviation (SD). Oligonucleotide loading was quantified using flow cytometry and thermogravimetric analysis (TGA) in a dynamic oxygen atmosphere with a heating ramp of 250°C (5 min) to 1000°C at a rate of 10°C/min. A Pfeiffer Vacuum ThermoStar mass spectrometer was used to measure MS signals: m/z = 18, m/z = 46, m/z = 44, and m/z = 15. Prodrug release was performed *in vitro* by stirring FdU_10_@SiO2 particles in PBS for 72 hours at 37°C. Samples were extracted at various time points and analyzed using fluorescence spectroscopy with an Edinburgh Inst. FLSP920 spectrofluorometer. The FdU_10_Cy5 release in PBS was fitted to a Korsmeyer–Peppas kinetic model.

### In cellulo studies

Murine malignant melanoma B16-F10 cells (Innoprot) were cultured in Iscove’s Modified Dulbecco’s Medium (IMDM, Panbiotech) supplemented with 10% fetal bovine serum (Fisher Scientific; Waltham, MA, USA) and antibiotics. FdU_10_Cy5@SiO_2_ particles were resuspended in the medium at 100 µg NPs/ml. At 2-, 24-, and 72-hours post-addition, cells were washed twice with PBS, fixed in 4% paraformaldehyde, stained, mounted, and analyzed using a Nikon AIR confocal microscope or flow cytometry with a CytoFLEX8® (Beckman Coulter) system. Cell cycle analysis was performed on cultures treated with T_10_@SiO_2_ and FdU_10_Cy5@SiO_2_ particles. At 24, 48, and 72 hours, cells were harvested, washed, fixed in 70% cold EtOH, and stained with Propidium Iodide. Immunofluorescence for Ki67 (Abcam) and p21 (Abcam) was conducted on cells after 72 h treatment (100 µg NPs / mL) and fixed with 4% paraformaldehyde 72 hours post-treatment, using anti-Ki67 and anti-p21 polyclonal antibodies, and visualized with Alexa Fluor 488 and Cy3 conjugated secondary antibodies (Invitrogen). All images were pseudocolored.

### Preclinical animal models

*In vivo* experiments were designed to minimize animal use, with ethical approval obtained from the Gobierno de Cantabria, Consejería de Medio Rural, Pesca y Alimentación (accreditation reference: PI-05-23). C57BL/6 mice (8 weeks old) were purchased from Janvier Labs. Animals were maintained, handled, and sacrificed following directive 2010/63/UE. Local *in vivo* treatments involved intra-scapular subcutaneous transplanting CD1 neonate mice with 1 × 10^5^ B16-F10 cells resuspended in 10 μl of IMDM containing antibiotics as previously described.^[26]^

After 7 days, mice were randomly divided into four groups receiving PBS, T_10_@SiO_2_, free FdU_10_Cy5, or FdU_10_@SiO_2_ at 300 µg FdU_10_Cy5/kg. Mice were euthanized 3 days post-treatment, and the tumors were weighed and evaluated in the IVIS® Biofluorescence equipment to quantify the Cy5 labeled prodrug using excitation/emission wavelengths of 650/670 nm during a 1-min exposure. The fluorescence intensity was quantified as the mean radiant efficiency (photons s⋅cm^-2^⋅sr)/(μW⋅cm^2^). Confocal microscopy imaging when used for quantification, was conducted with identical settings for both control and experimental samples, utilizing a Nikon AIR microscope.

Systemic *in vivo* treatments involved intra-scapular subcutaneous transplanting with 2 × 10^5^ B16-F10 cells resuspended in 10 μl of IMDM-containing antibiotics as previously described.^[26]^ For the treatment phase, mice were randomly divided into 4 groups, each consisting of 10 mice. These groups received PBS, FdU_10_@SiO_2_, FdU_10_@SiO_2_@GFP:VEGFRbp, FdU_10_@SiO_2_@PA17. Mice were treated 3 times on days 4, 7 and 10 after cell transplant at a concentration of 120 μg/kg of FdU_10_-Cy5 /dose. The mice were euthanized 12 days after the transplant and their tissues were collected and fixed in formalin. Tumors were weighed and evaluated in the IVIS® Biofluorescence equipment to quantify the Cy5 labeled prodrug using excitation/emission wavelengths of 650/670 nm during a 1-min exposure. The fluorescence intensity was quantified as the mean radiant efficiency (photons s⋅cm^-2^⋅sr)/(μW⋅cm^2^). Confocal microscopy imaging was conducted with identical settings for both control and experimental samples, utilizing a Nikon AIR microscope. VEGF and TEM8 receptors were immunostained with anti-VEGF Receptor 1 (Thermofisher) and TEM8/ATR (Abcam), antibodies, and were visualized with an Alexa Fluor 488 conjugated secondary antibody (Invitrogen). Tumors were evaluated using an IVIS® Biofluorescence system and confocal microscopy using a Nikon AIR microscope. All images were pseudocolored.

### Design, production, purification, and functionalization of the ligand proteins

Recombinant gene constructs encoding GFP:VEGFRbp were cloned into pET 15b plasmid systems (Novagen) by General Biosystems, Inc. (Morrisville, USA). Both ligand proteins were fused to a 10-histidine tag at their amino-terminus (Figure S13) for electrostatic particle functionalization as in previous studies.^[52,55]^ Constructs were transformed and expressed in One Shot™ BL21 (DE3) *E. coli* (Thermo Fisher Scientific). Proteins were purified using Ni-TED columns (Protino® Ni-TED, Macherey-Nagel) following standard biochemical procedures. Protein analysis was conducted using SDS-PAGE with Coomassie-stained gels analyzed using the BioRad GelDoc EZ system software. FdU_10_@SiO_2_ functionalization with both ligands was based on a previous protocol of our group. In brief, 100 μg of particles were immersed in 500 μL PBS containing saturating amounts of the tagged protein (ca. 0.5 mg/mL) at room temperature. The mixture was sonicated in a water bath for 5 min at 4 °C. Unbound protein was removed by repeated centrifugation. SDS-PAGE electrophoresis was used to quantify the protein captured on the surfaces of the particles. These were stripped in Laemmli sample buffer (BioRad) at 90 °C. The stripped protein was loaded in precast Mini-Protean® TGX™, BioRad gels for SDS-PAGE analysis. The semi-quantification of the total amount of protein on the surface of the particles was performed on Coomasie-stained gels using the software of the BioRad GelDoc EZ system.

## Statistics

Results are expressed as mean values along with their corresponding standard deviations (SD). Statistical differences in means were assessed using a t-test or ANOVA, conducted through Graphpad Prism, and significance was established at p < 0.05. In cases where ANOVA yielded significant results, pairwise comparisons were carried out. The total number of events and the confidence levels achieved in the experiment (n) are all specified in each figure caption for reference.

## Supporting Information

Supporting Information is available from the Wiley Online Library or from the author.

## Author Contributions

ARV, AD, NN, AML, AA, and LGH, performed the experiments. All authors discussed the results and wrote the manuscript. MLF, RE, and CF obtained the funding. All authors have approved the final version of the manuscript.

## Acknowledgements

We acknowledge the financial support from the Spanish Instituto de Salud Carlos III, under Project ref. DTS24/000237, PI22/00030, and PI23/00261 co-funded by the European Regional Development Fund, “Investing in your future,” Grant TED2021-129248 BeI00 the Spanish Ministerio de Ciencia e Innovación (MICINN) Projects PID2020-118145RB-I00 and TED2021-129248B-100, co-funded by the European Union FEDER funds; the Gobierno Regional de Cantabria and IDIVAL for the project Refs INNVAL21/19, NEXTVAL 22/12, and AR IDI-020-022 fellowship; N.N. held a predoctoral contract grant (PRE2021-097856). We also thank Oligonucleotide synthesis was performed by the ICTS ‘‘NANBIOSIS” (CIBER BBN) and specifically by the https://www.nanbiosis.es/portfolio/u29-oligonucleotide-synthesis-platform-osp/ oligonucleotide synthesis platform (OSP) U29 at IQAC-CSIC.

We gratefully acknowledge Ms. Débora Muñoz and Claudia Zamarrón for their technical help, Drs. Iñigo Casafont and Juan Carlos Acosta for their assistance with the senescence experiments, and Dr. Rafel Valiente for his help with fluorimetry experiments. The figures and graphs have been created with BioRender software (BioRender.com, License ID: 9519A1C8-0002). We would like to express our gratitude to the COST Action DARTER 17103 for providing a collaborative environment that facilitated the development of this research.

## Data Availability Statement

The data presented in this study are available on request from the corresponding author.

---

This research introduces a novel approach for treating malignant melanoma using a triple-strategy therapy. A Floxuridine-based oligonucleotide prodrug is encapsulated in silica nanoparticles, targeting tumor neovasculature. Results from *in vitro* and preclinical models show significant tumor reduction, highlighting the potential of this targeted delivery system in improving treatment outcomes for metastatic melanoma.

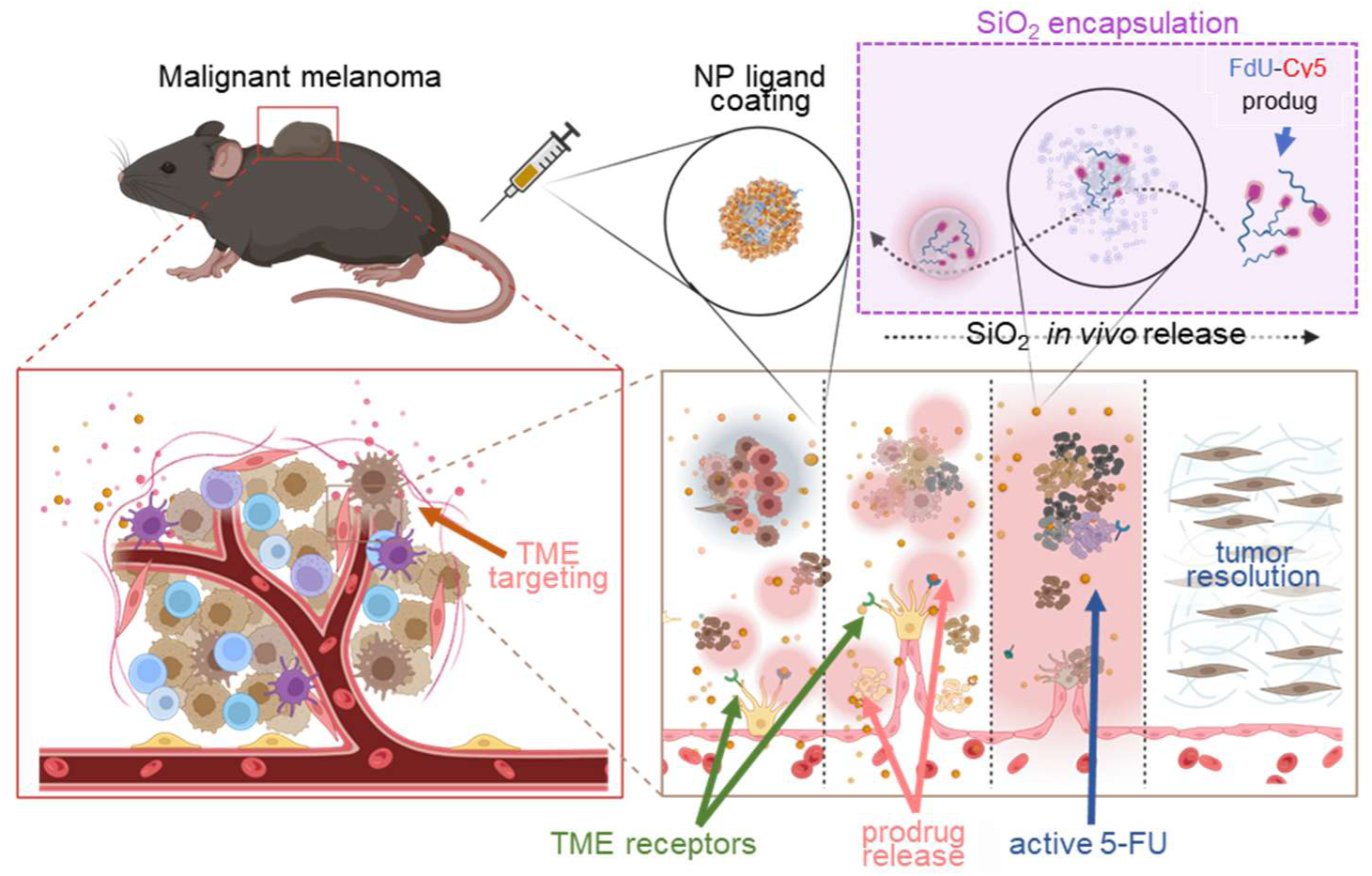

