## Supplementary Information for "Targeted Tumor Microenvironment Delivery of Floxuridine Prodrug *via* Soluble Silica Nanoparticles in Malignant Melanoma as a Model for Aggressive Cancer Treatment"

**Table of contents:**


- **Supplementary methods**:
  - Oligonucleotide synthesis.
  - Oligonucleotide characterization.
- **Supplementary tables:**
  - Table S1. Synthesis and characterization of formulations #01 to #04 $FdU_{10}Cy5@SiO2$ particles.
  - Table S2. Fitting of $FdU_{10}Cy5$ release from $FdU_{10}@SiO_2$ particles to Karls-Peppas-Meyer model.
- **Supplementary figures:**
  - Figure S1. HPLC profile and MALDI-TOF of $FdU_{10}Cy5$ and $T_{10}Cy5$.
  - Figure S2. Characterization of Nanoparticles from Different Formulations.
  - Figure S3. Relative embedment of $FdU_{10}Cy5$ within $FdU_{10}Cy5@SiO_2$ nanoparticles.
  - Figure S4. Characterization of control $T_{10}Cy5@SiO_2$ nanoparticles.
  - Figure S5. Thermogravimetric Analysis (TGA) of the loaded nanoparticles.
  - Figure S6. Fluorimetry Measurement of $FdU_{10}$-Cy5 oligonucleotide release
  - Figure S7. Cell uptake of $FdU_{10}$-Cy5 oligonucleotide.
  - Figure S8. Subcellular localization of the prodrug.
  - Figure S9. Raw flow cytometry cell cycle data.
  - Figure S10. Senescence assessment in treated cells.
  - Figure S11. IVIS® Cy5 Bioimaging of tumors treated intratumorally.
  - Figure S12. Amino acid sequences of the ligand proteins.
  - Figure S13. Confocal microscopy of the tumor live immunostained for TEM8.
  - Figure S14. SDS-PAGE analysis of the recombinant proteins used.
  - Figure S15. Changes in body weight of mice during the period of systemic treatment.
  - Figure S16. IVIS® Cy5 Bioimaging of tumors after systemic treatment.
  - Figure S17. Antitumoral effect of the targeted Nanoparticles.
  - Figure S18. Intratumoral Neovasculature Analysis Following Systemic Treatments

**Supplementary methods**

*Oligonucleotide synthesis*: The oligonucleotide synthesis was performed on an H-8 DNA synthesizer (K&A Laboratories, Germany) at a 1 μmol scale using phosphoramidite solid-phase protocols. The support used was the commercial DMTCy5-CPG (LGC-Bioresearch, Germany). The strands were ensembled using the corresponding 5'-DMT-3'-[(2-cyanoethyl)-(N,N-diisopropyl)]-phosphoramidites at 0.1 M concentration in anhydrous acetonitrile. Cleavage from the solid support was done treating the sample with a 32 % aqueous ammonia solution at 55 ºC for 1 hour. Then, the oligonucleotides were desalted using Sephadex G-25 columns and used without any additional purification.

*Oligonucleotide characterization*: The purity of the oligonucleotides was analyzed by HPLC and mass spectroscopy (MALDI-TOF). Solvents for HPLC were prepared using triethylammonium acetate buffer and acetonitrile. Buffer A was 5% acetonitrile in 0,1 M of triethylammonium acetate and Buffer B was 70% acetonitrile in 0,1 M of triethylammonium acetate. HPLC was performed on a Waters chromatography system with a Waters 2996 Photodiode Array Detector using a Nucleosil column 120 C18 (250x4mm). The prepared oligonucleotides were analyzed with a gradient of 0-50% solution B in 20 minutes. Mass spectra were recorded on a MALDI Voyager time-of-flight spectrometer using 2',4',6'-trihydroxyacetophenone monohydrate and ammonium citrate dibasic as matrices.

### Supplementary tables

**Table S1.** Synthesis and characterization of formulations #01 to #04 $FdU_{10}Cy5@SiO_2$ particles

| Name | Synthesis | | | | | Characterization | |
|---|---|---|---|---|---|---|---|
| | **EtOH/M** | **$NH_3$ [M]** | **TEOS [M]** | **$H_2O$ [M]** | **$[FdU_{10}]$ [M] (µg FdU10/mg $SiO_2$)** | **Z-size [nm]** | **Z-pot [mV]** |
| $FdU_{10}@SiO_2$ #01 | 10.0 | 1.22 | 0.24 | 9.99 | $1.4 \times 10^{-5}$ (4.5 µg) | 499 ± 23 | -47 ± 1 |
| $FdU_{10}@SiO_2$ #02 | 12.5 | 0.41 | 0.12 | 5.00 | $0.7 \times 10^{-5}$ (4.5 µg) | 343 ± 20 | -33 ± 7 |
| $FdU_{10}@SiO_2$ #03 | 13.7 | 0.34 | 0.08 | 3.33 | $0.46 \times 10^{-5}$ (4.5 µg) | 244 ± 15 | -31 ± 3 |
| $FdU_{10}@SiO_2$ #04 | 14.3 | 0.31 | 0.06 | 2.5 | $1.6 \times 10^{-6}$ (4.5 µg) | 209 ± 2 | -28 ± 2 |

**Table S2.** Fitting of $FdU_{10}Cy5$ release from $FdU_{10}@SiO_2$ particles to Karls-Peppas-Meyer model.

| Time / h | $FdU_{10}Cy5$ release (%) | Fit KMP model (%) | SD |
|---|---|---|---|
| 0 | 0 | 0 | 0 |
| 2 | 21 | 31 | 85 |
| 4 | 34 | 34 | 0 |
| 6 | 43 | 36 | 43 |
| 24 | 48 | 44 | 13 |
| 48 | 51 | 49 | 4 |
| 72 | 53 | 52 | 0 |
| 144 | 54 | 58 | 18 |

| | |
|---|---|
| **K** | 27.66 |
| ***n*** | 0,148 |

| | |
|---|---|
| **SSD** | 162.46 |

**Supplementary figures**

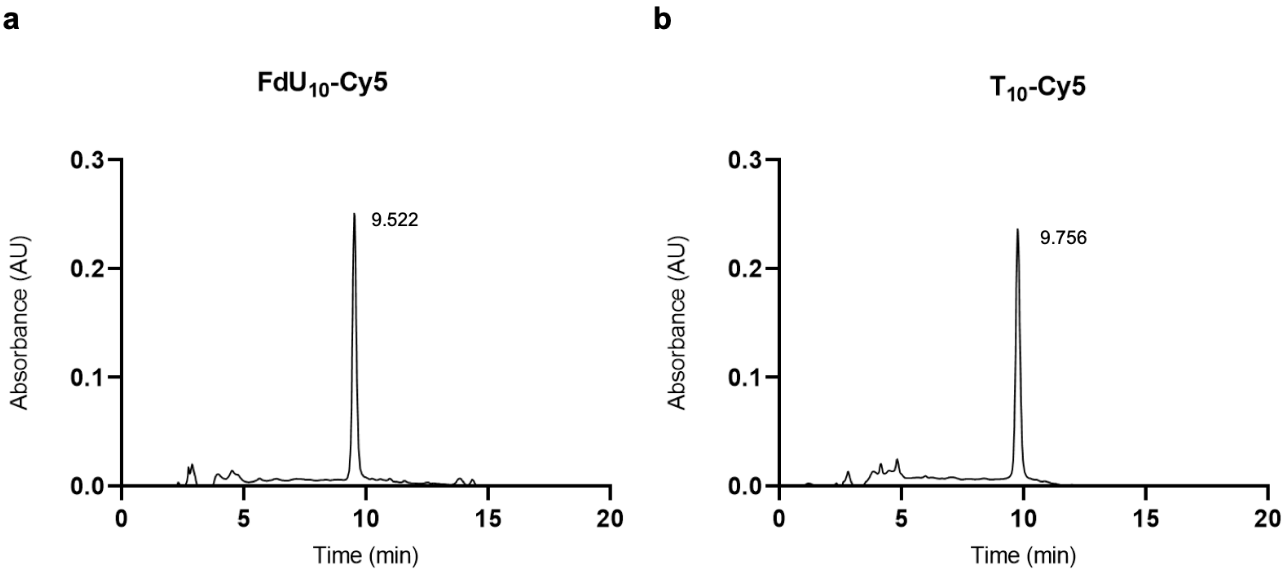


**Figure S1A. a.** HPLC profile of **a** $FdU_{10}Cy5$ and **b** $T_{10}Cy5$.

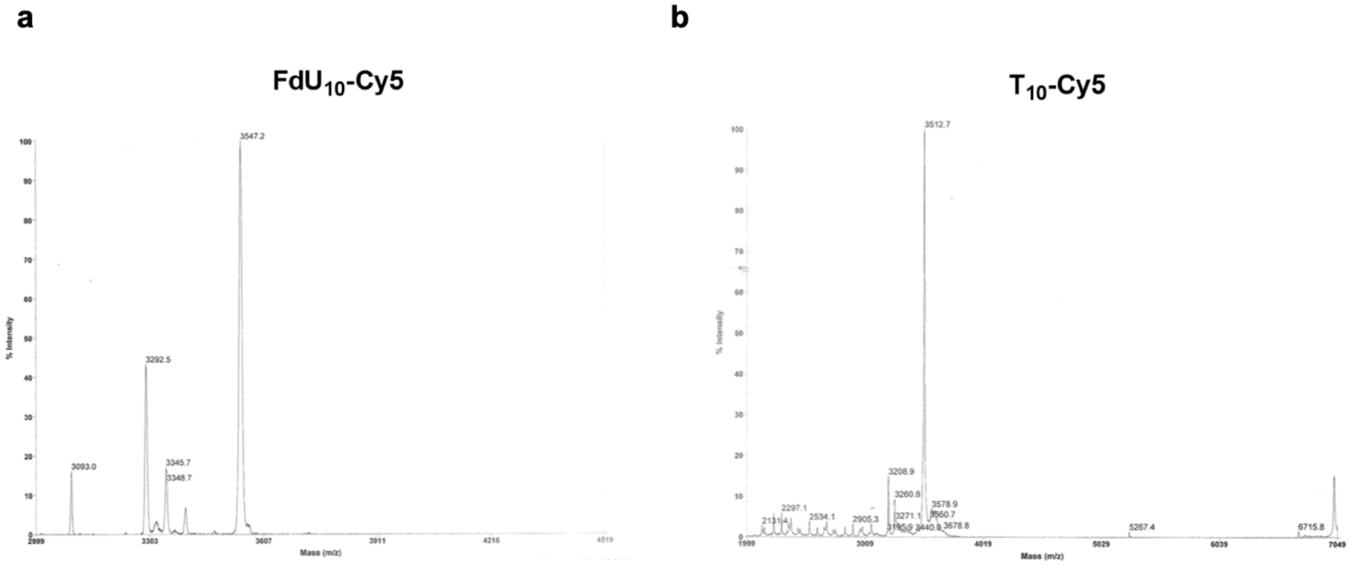


**Figure S1B.** MALDI-TOF of **a** $FdU_{10}Cy5$. Expected molecular weight: 3550.8 g/mol; found molecular weight: 3547.2 g/mol and **b** $T_{10}Cy5$. Expected molecular weight: 3511.8 g/mol; found molecular weight: 3512.7 g/mol.

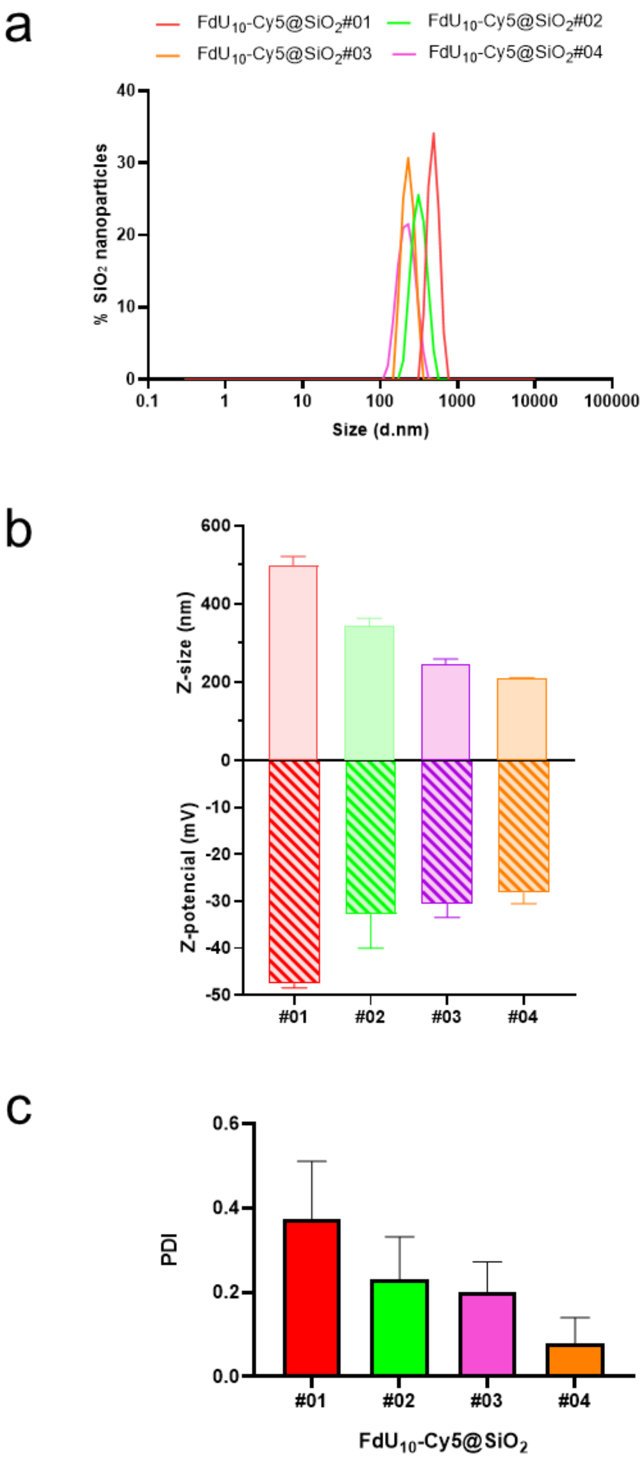


**Figure S2. Characterization of Nanoparticles from Different Formulations**. **A**. Size distribution profile of $FdU_{10}Cy5@SiO_2$ #01 to #04 particles by DLS. **B.** Hydrodynamic diameter (Z-size /nm) and Z-potential (mV) of $FdU_{10}Cy5@SiO_2$ #01 to #04 **C**. Polydispersity Index (PDI) of particles $FdU_{10}Cy5@SiO_2$ #01-04 measured by DLS.

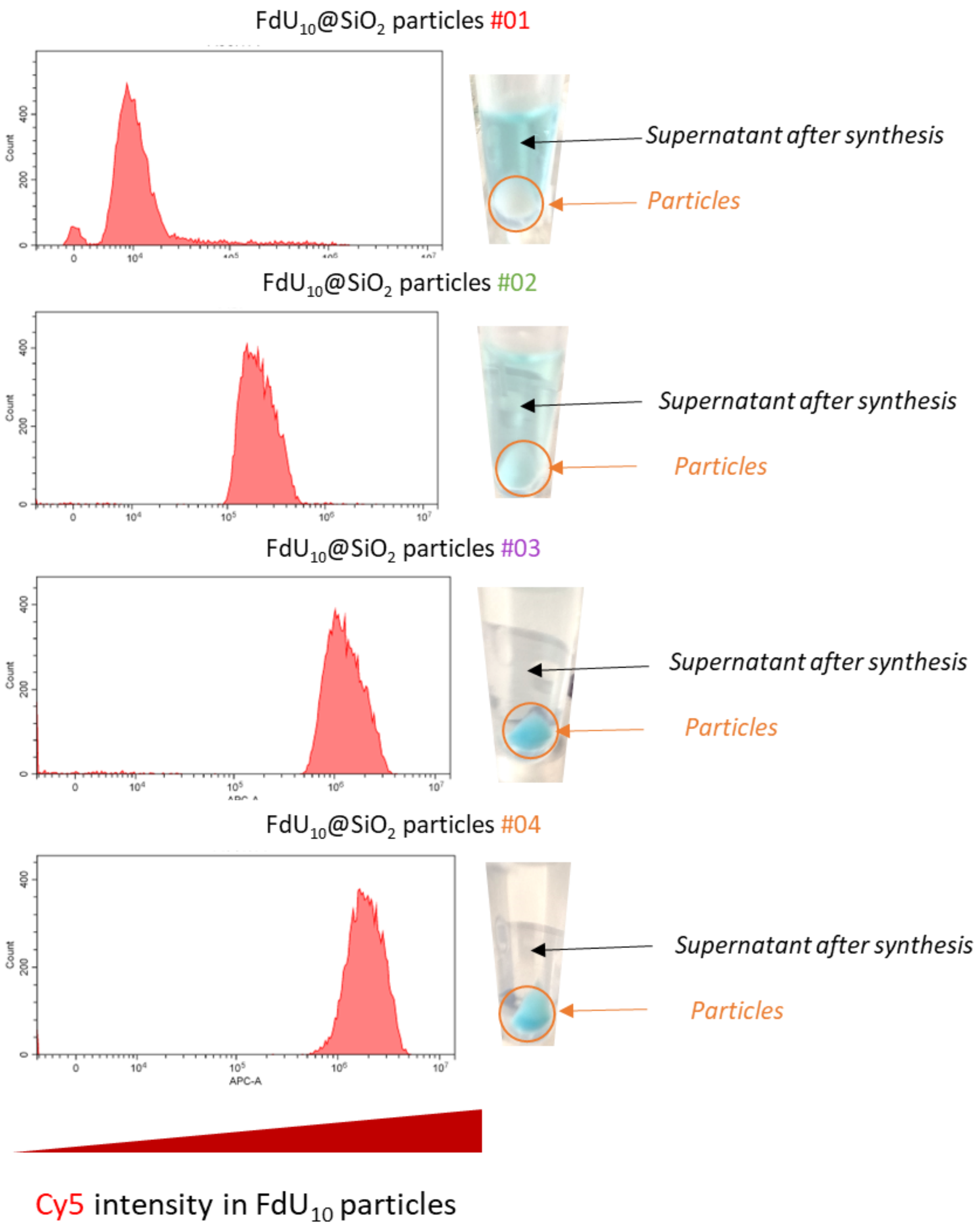


**Figure S3.** Relative embedment of $FdU_{10}Cy5$ within $FdU_{10}Cy5@SiO_2$ nanoparticles measured by flow cytometry. The blue color in the nanoparticle pellet or supernatant shows the localization of the Cy5 fluorophore attached to the prodrug.

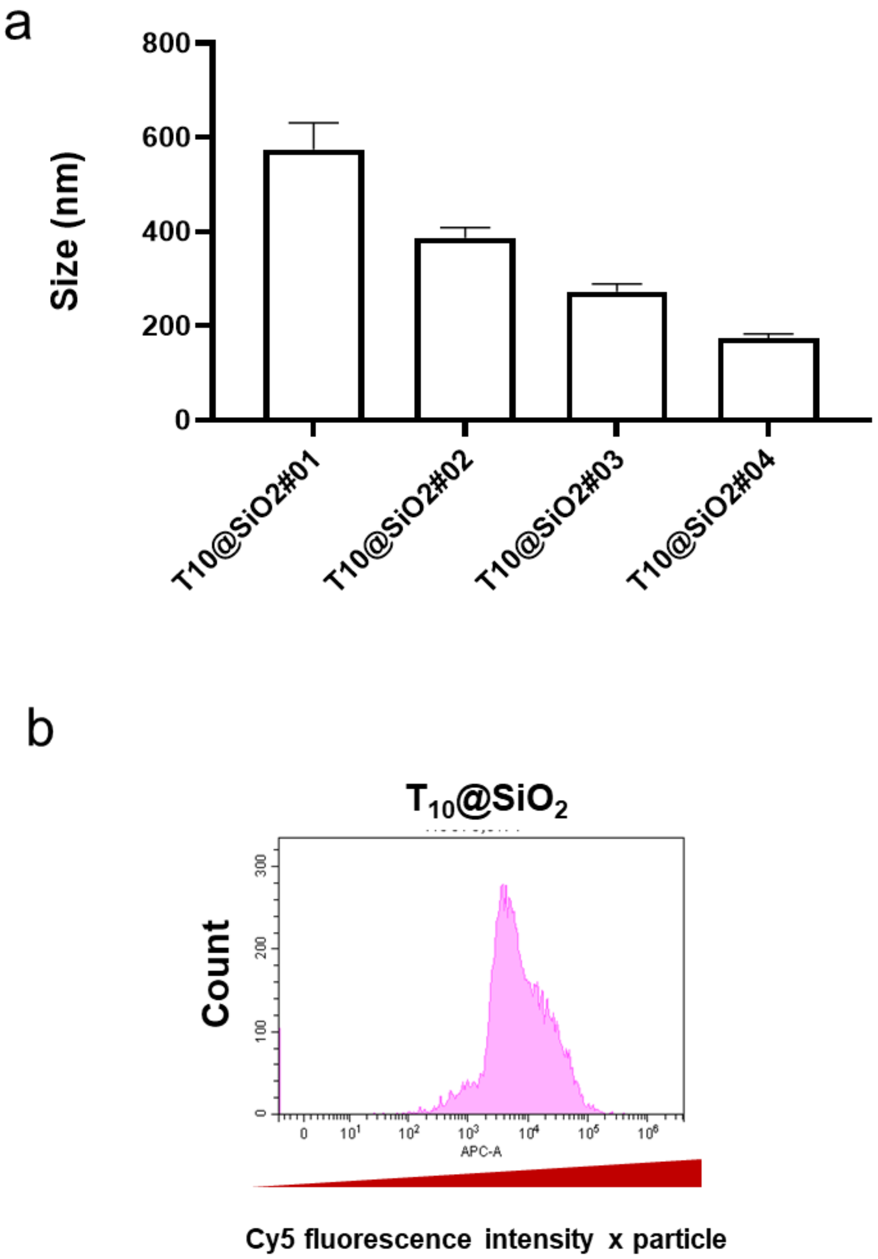


**Figure S4. Characterization of Control $T_{10}$Cy5@$SiO_2$ Nanoparticles. A.** Size measured by DLS of the control $T_{10}$Cy5@$SiO_2$ particles produced following synthesis #01 to #04 protocols. **B.** Relative embedment of $T_{10}$Cy5 oligo within $T_{10}$@$SiO_2$#04 nanoparticles measured by flow cytometry.

**Control $SiO_2$ #04 particles (without $FdU_{10}$-Cy5)**

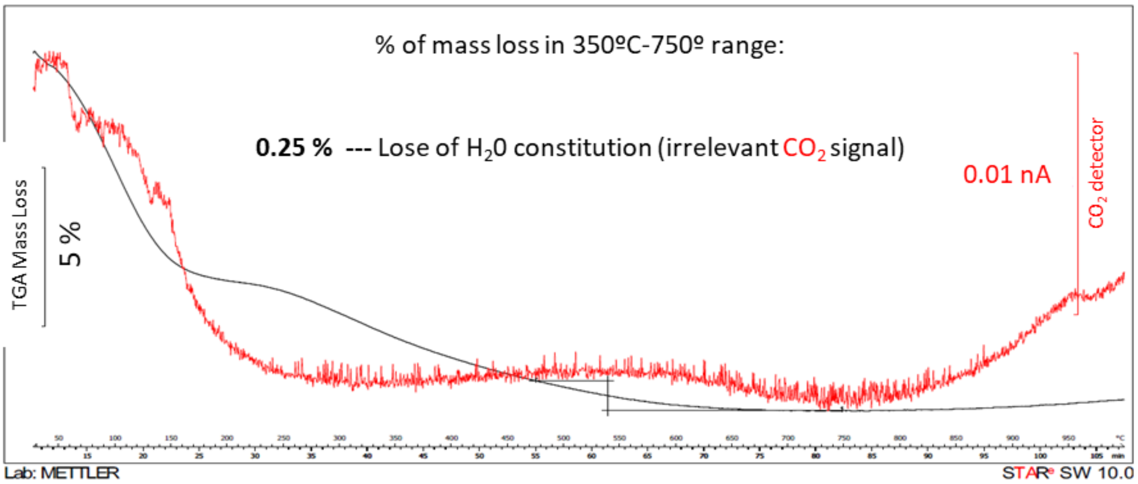


**$FdU_{10}$-Cy5@$SiO_2$ #04 particles**

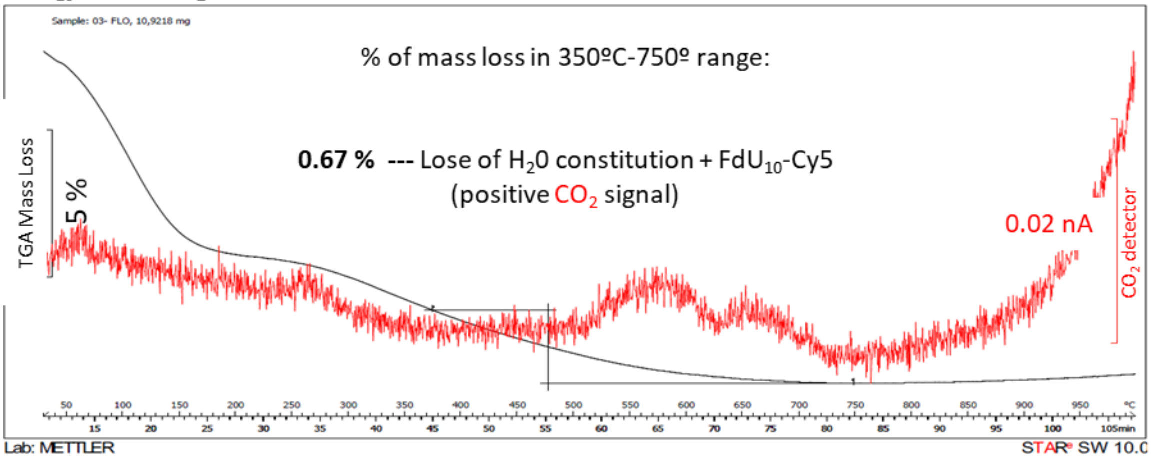


**% of loaded $FdU_{10}$-Cy5= 0.67%-0.25%=0.44 %**
(mass FdU10/total mass of particle)

**% encapsulation efficiency = 97.1 %**
(mass of entrapped $FdU_{10}$-Cy5/initial mass of $FdU_{10}$-Cy5)

**Figure S5. Thermogravimetric Analysis (TGA) of the Loaded Nanoparticles.** TGA diagram of control (empty) $SiO_2$ particles and $FdU_{10}Cy5@SiO_2$ particles ranging from 0 ºC to 1000 ºC, with an increase of 5 ºC / min. The black line corresponds to mass loss, and the red line corresponds to the $CO_2$ detector signal. The temperature range of 350°C - 750°C indicates the decomposition of organic mass directly related to the $FdU_{10}Cy5$ oligonucleotide.

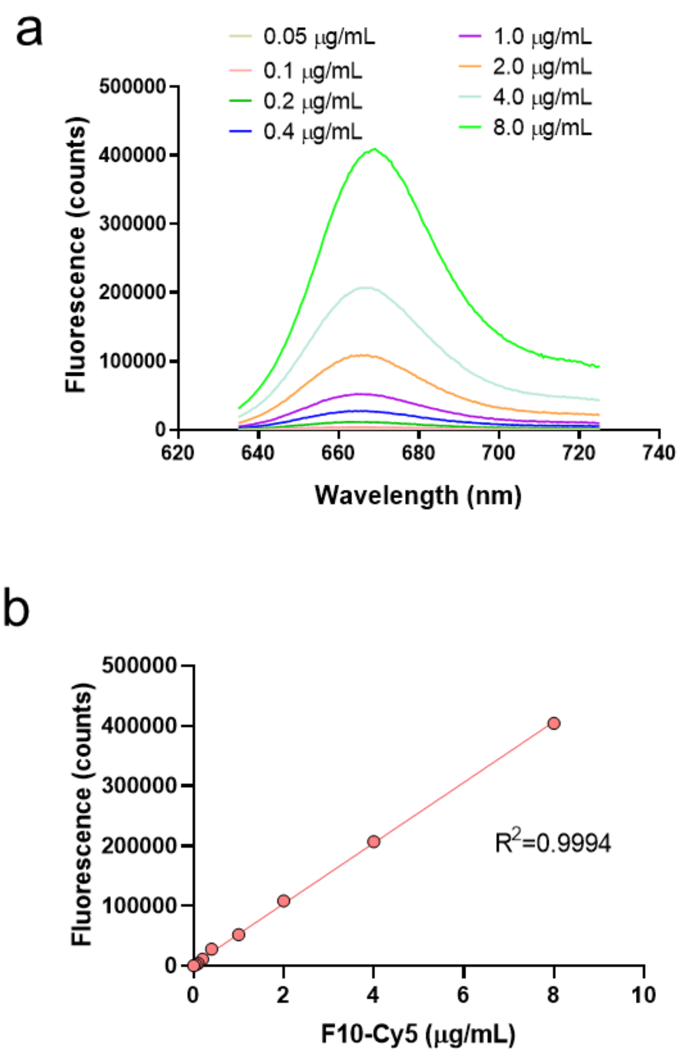


**Figure S6. Fluorimetry Measurement of $FdU_{10}Cy5$ Release. A.** Measurement of $FdU_{10}Cy5$ oligonucleotide release in PBS 1x solution. **B**. Calibration curve for direct quantification of $FdU_{10}Cy5$ during the *in vitro* release study.

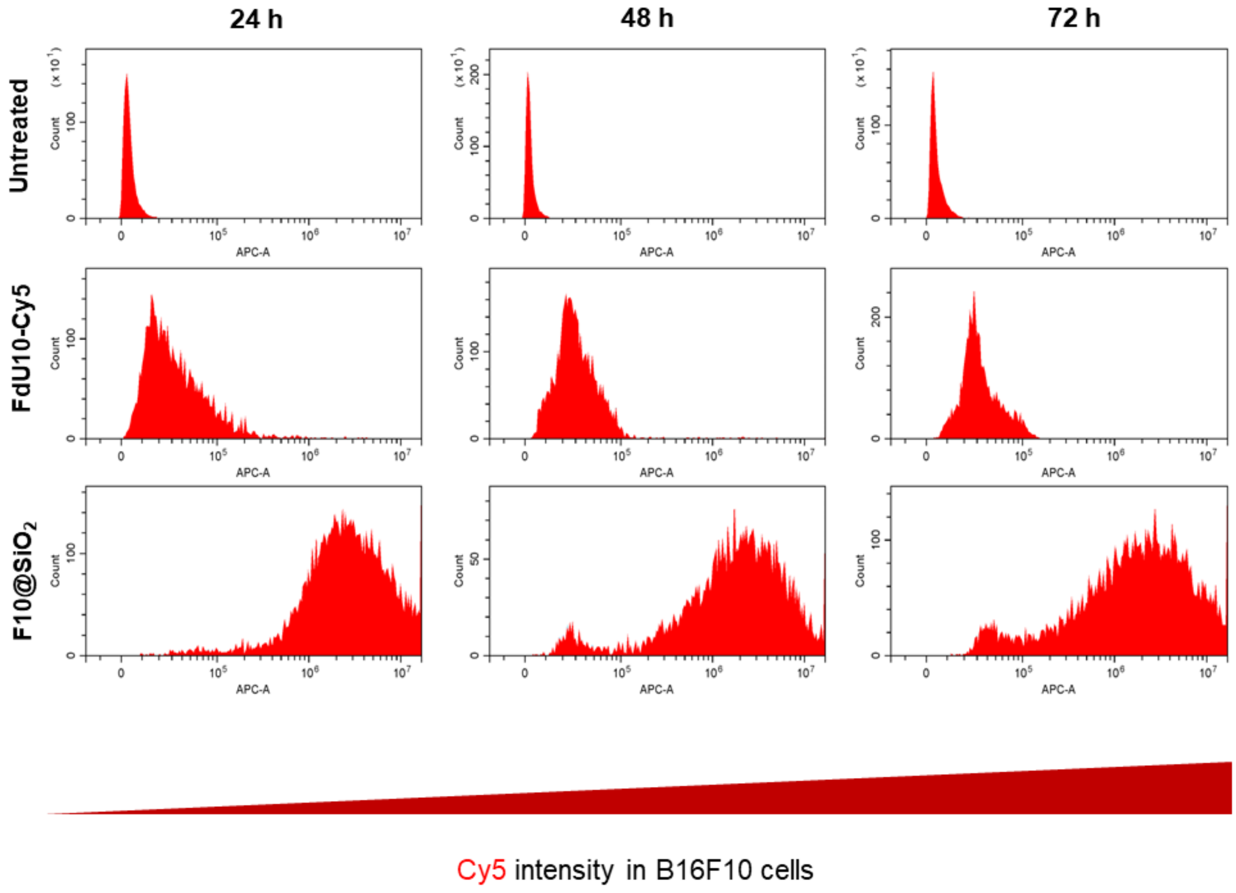


**Figure S7. Cell Uptake of $FdU_{10}$ Oligonucleotide.** Flow cytometry histograms illustrating the cellular uptake of $FdU_{10}Cy5$ oligonucleotide by melanoma cells at 24 hours, 48 hours, and 72 hours post-treatment.

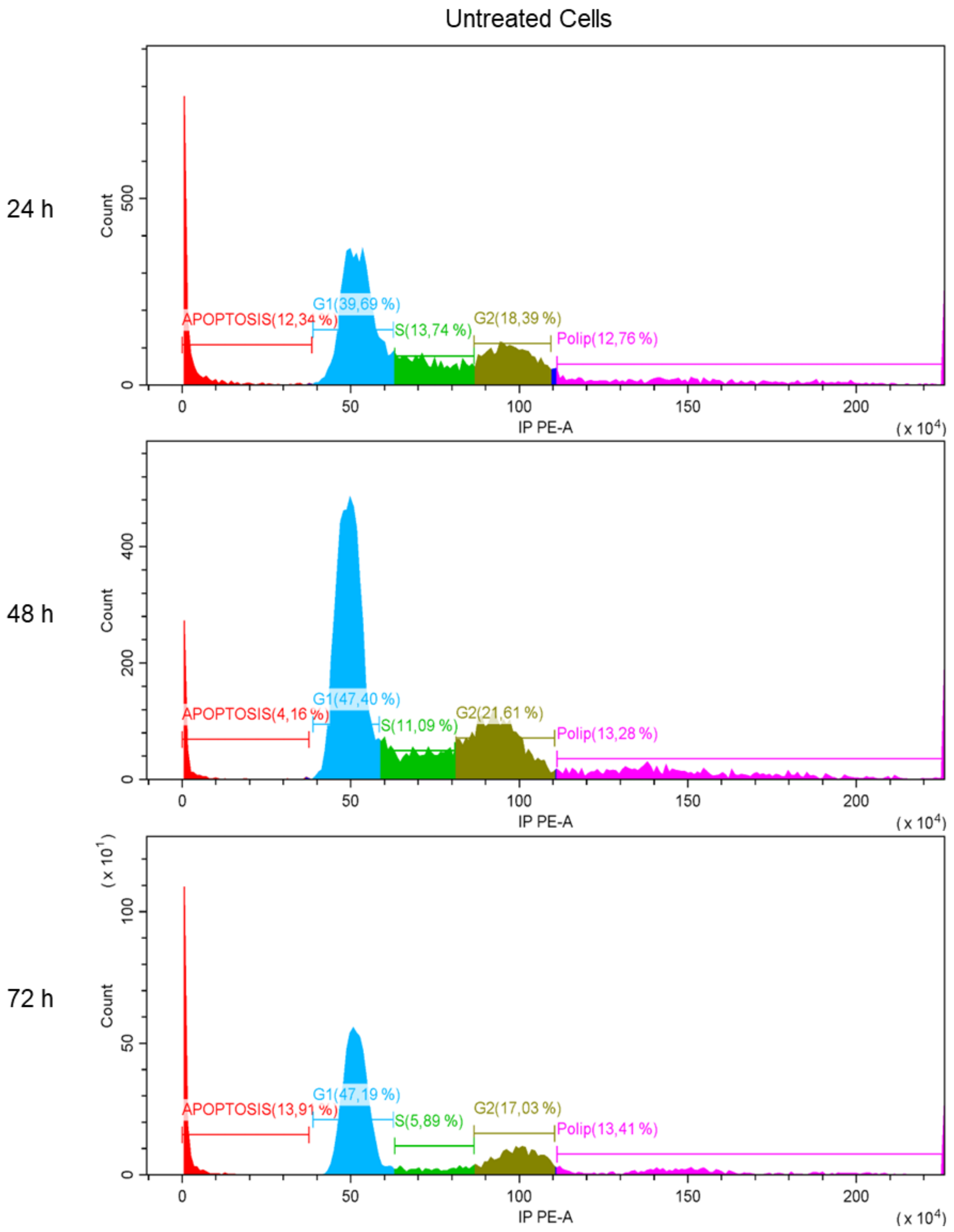


**Figure S8A. Raw Flow Cytometry Cell Cycle Data:** Histograms of untreated cell cycle analyses using propidium iodide (PI).

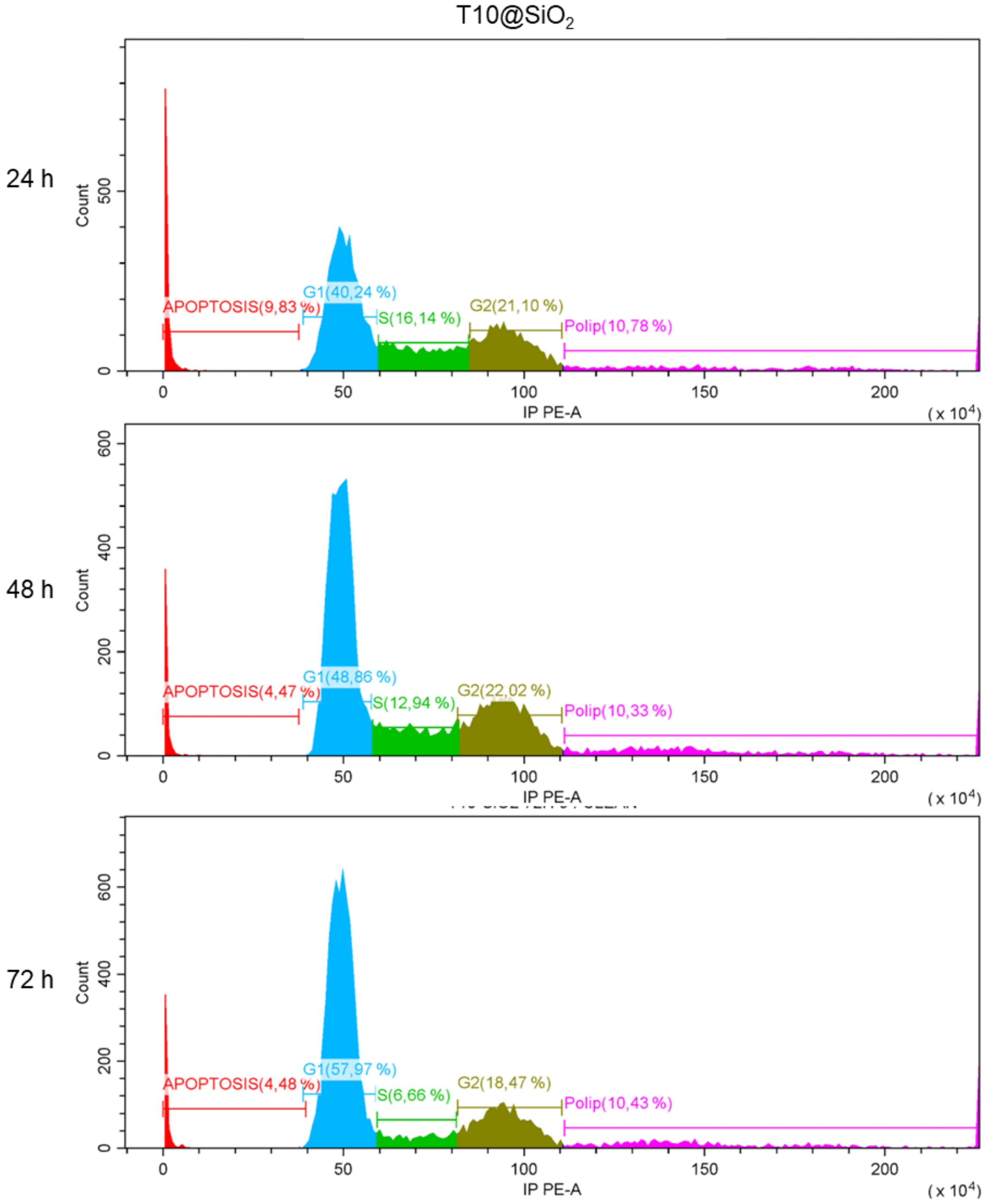


**Figure S8B. Raw Flow Cytometry Cell Cycle Data:** Histograms of cell cycle analyses using propidium iodide (PI) in melanoma cells treated with control $T_{10}@SiO_2$ nanoparticles.

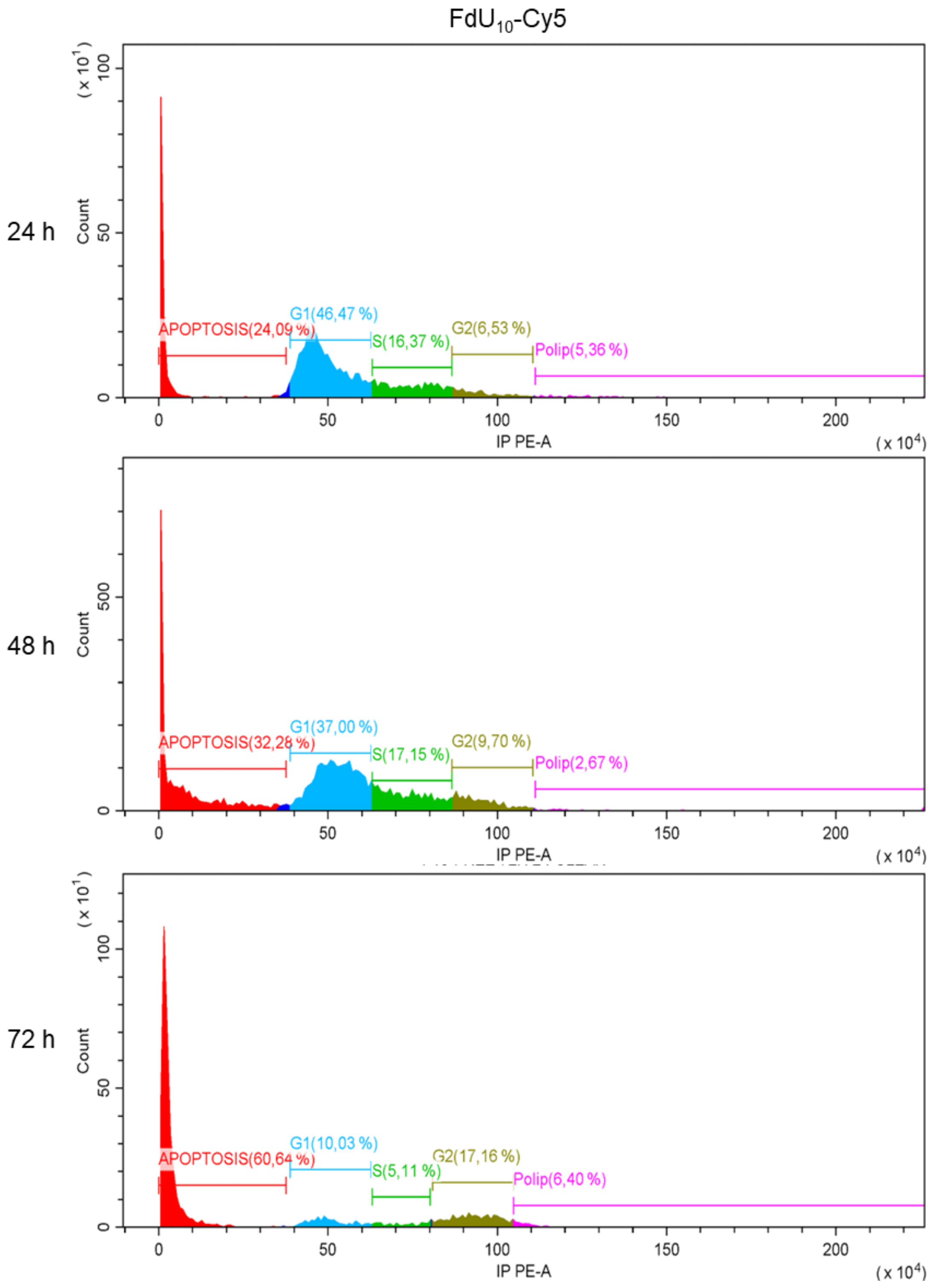


**Figure S8C. Raw Flow Cytometry Cell Cycle Data:** Histograms of cell cycle analyses using propidium iodide (PI) in melanoma cells treated with $FdU_{10}Cy5$ prodrug.

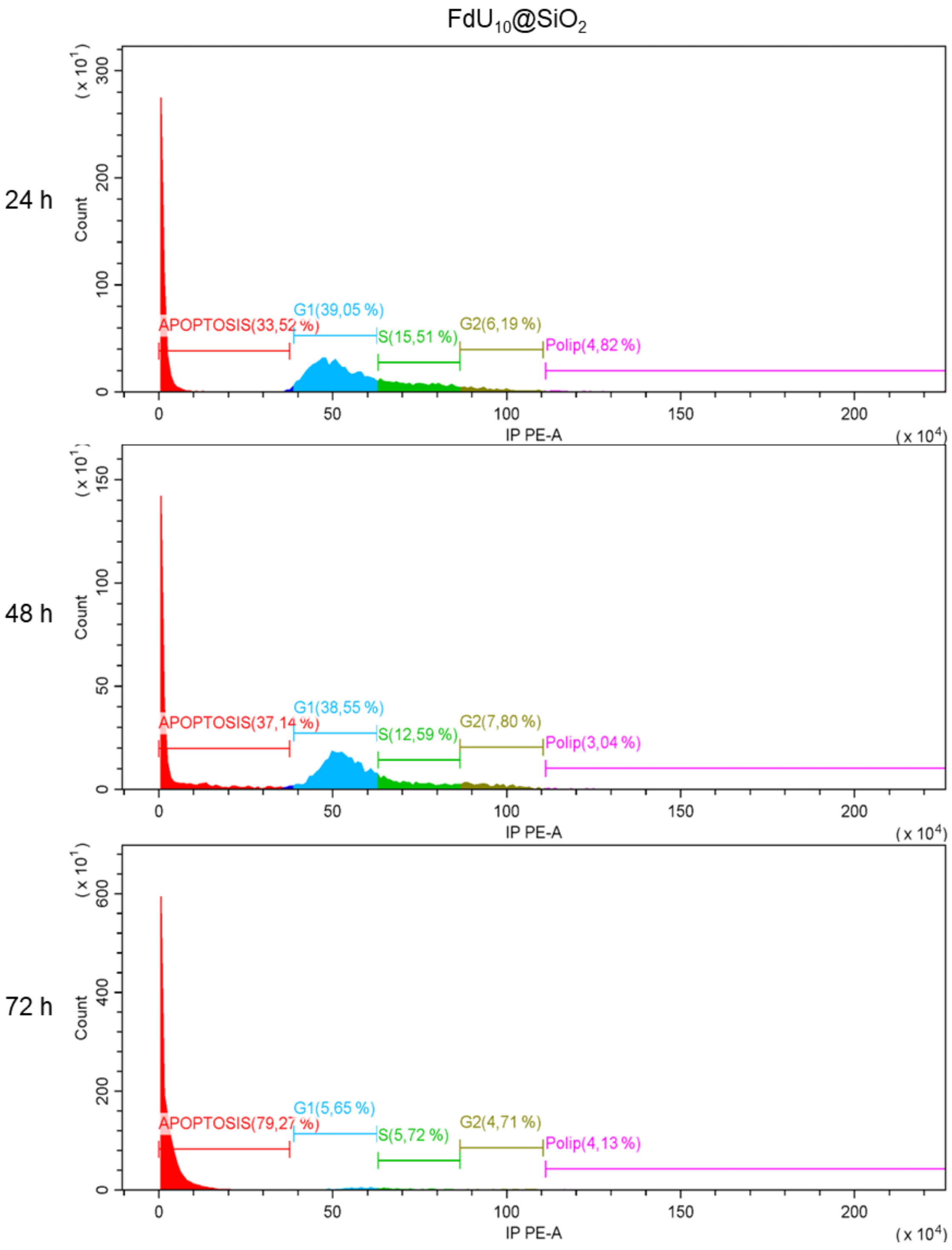


**Figure S8D. Raw Flow Cytometry Cell Cycle Data:** Histograms of cell cycle analyses using propidium iodide (PI) in melanoma cells treated with $FdU_{10}@SiO_2$ nanoparticles.

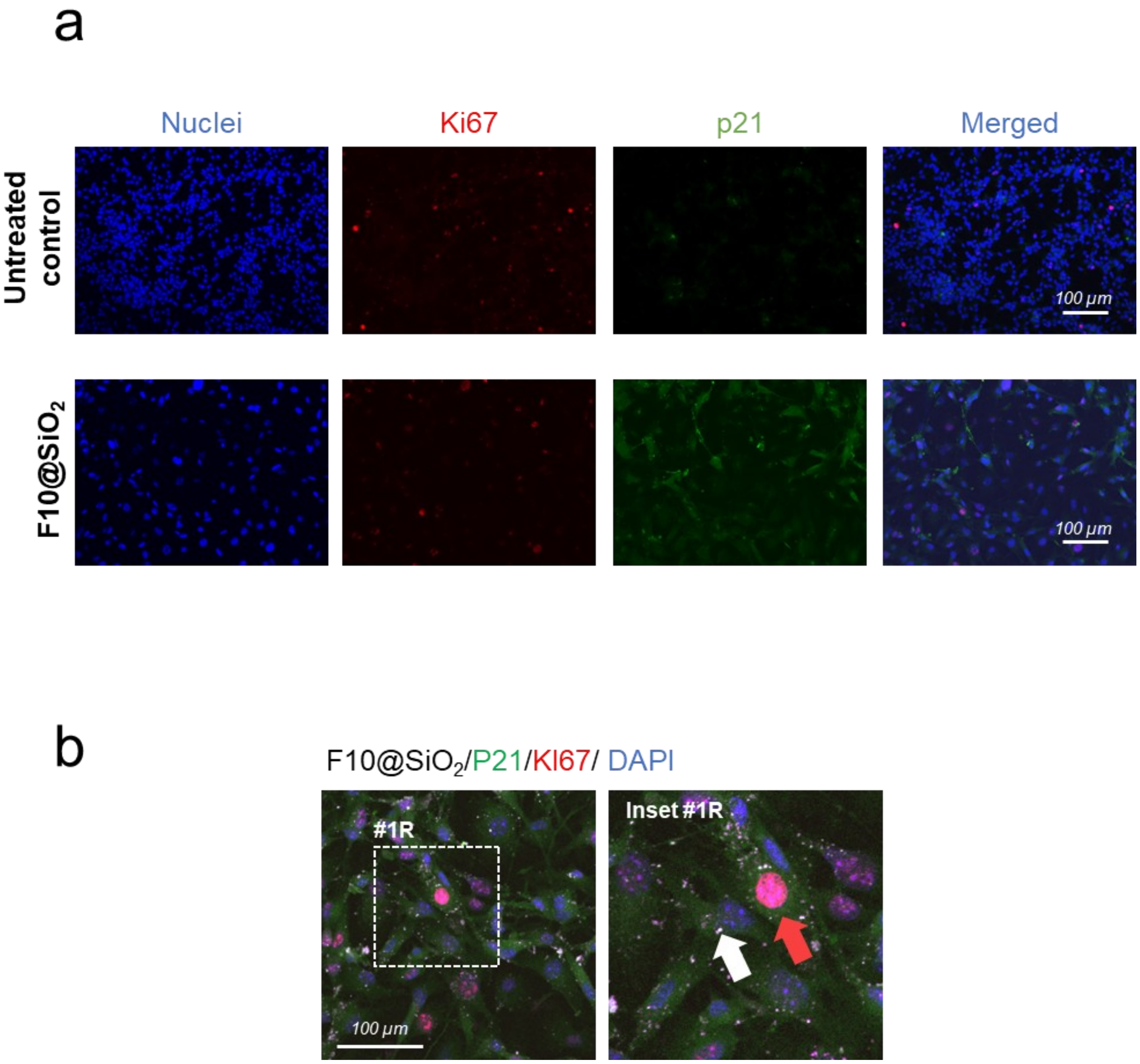


**Figure S9. Senescence Assessment in Treated Cells. A.** Confocal microscopy images of untreated melanoma cells and $FdU_{10}@SiO_2$ treated after 72 h. Comparative analysis of decreased Ki67 expression (proliferative marker) and increased p21 expression (senescent marker) in treated samples **B.** Confocal microscopy image of melanoma cells immunostained for p21 (green channel) and Ki67 (red channel) with intracellular $FdU_{10}@SiO_2$ nanoparticles (white channel, arrow) incubated with cells for 72 hours. Cell nuclei are stained with DAPI (blue channel). Some nuclei are positively immunostained for Ki67 (red arrow).

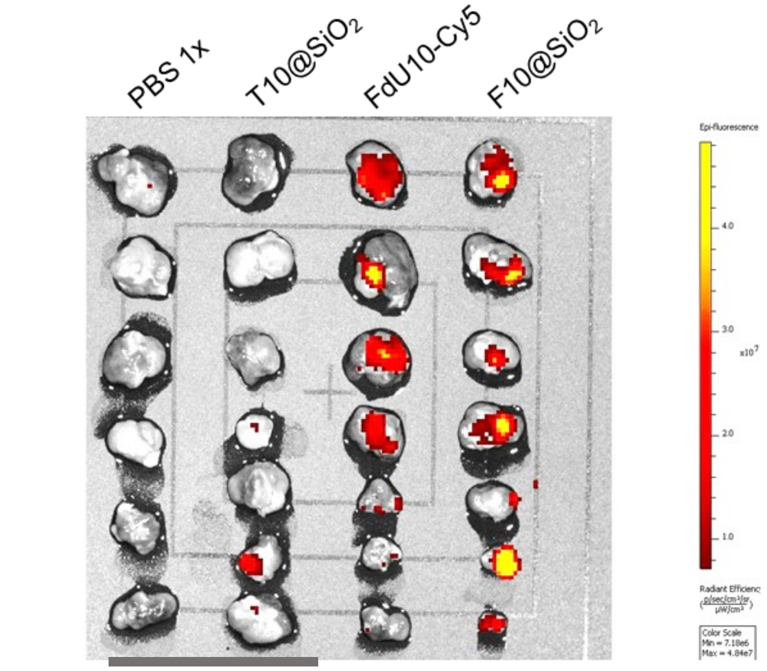


**Figure S10. IVIS® Cy5 Bioimaging of Tumors Treated Intratumorally**. Malignant melanoma tumors were collected postmortem from mice 72 hours after receiving the indicated intratumoral injections. The fluorescence was documented using an IVIS® biofluorescence imaging system. The fluorescence displayed corresponds to oligonucleotides labeled with Cy5, highlighting the accumulation of the treatments within the tumors. Scale bar (grey)=50 mm

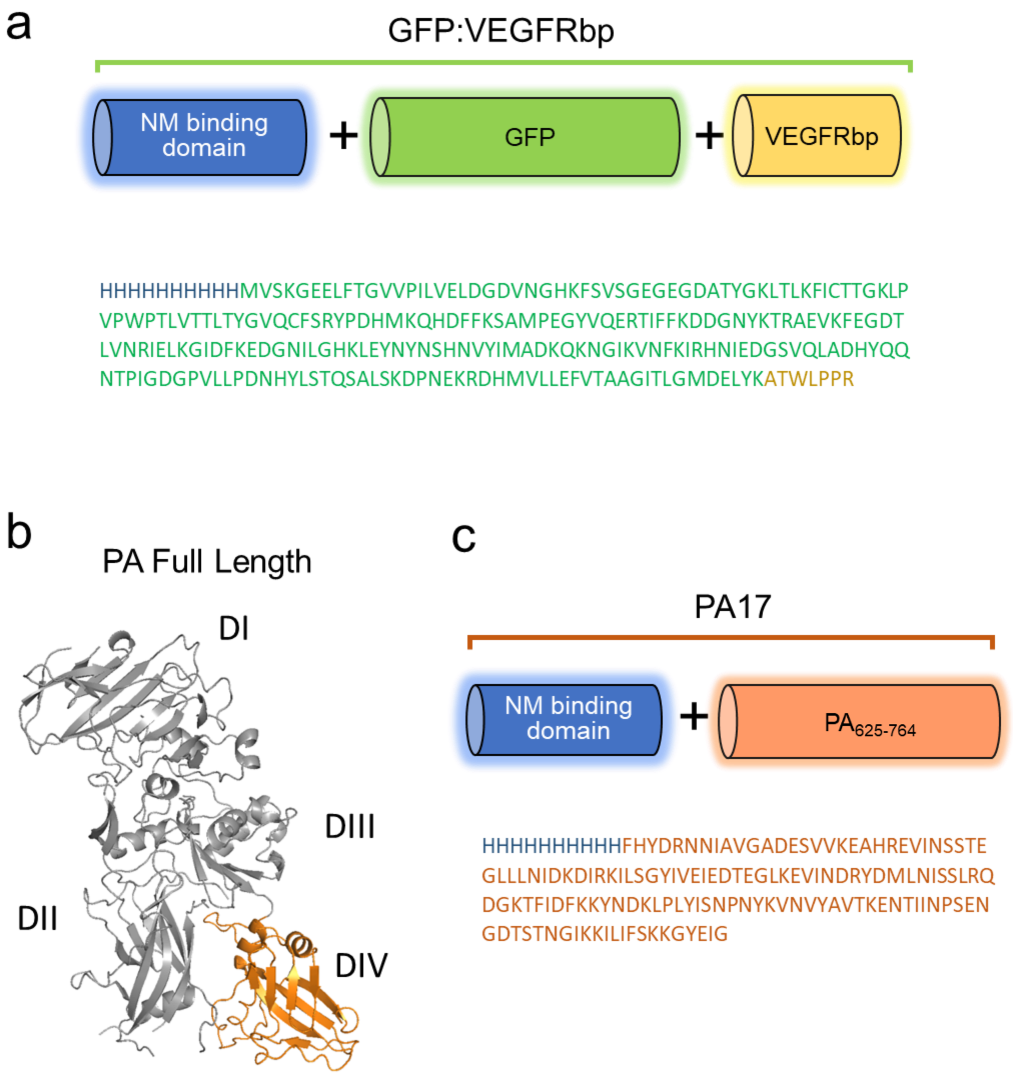


**Figure S11. Amino Acid Sequences of the Ligand Proteins.** Different parts of the proteins are highlighted in different colors for clarity. The 10xHis polycationic sequence (blue color) acts as the nanomaterial binding domain. a. Sequence of the GFP:VEGFRbp protein. b. The structure of the PA protein. The various domains of PA, with a focus on Domain IV (DIV), are indicated. c. Design of the engineered PA17 protein-ligand, showing the functional components.

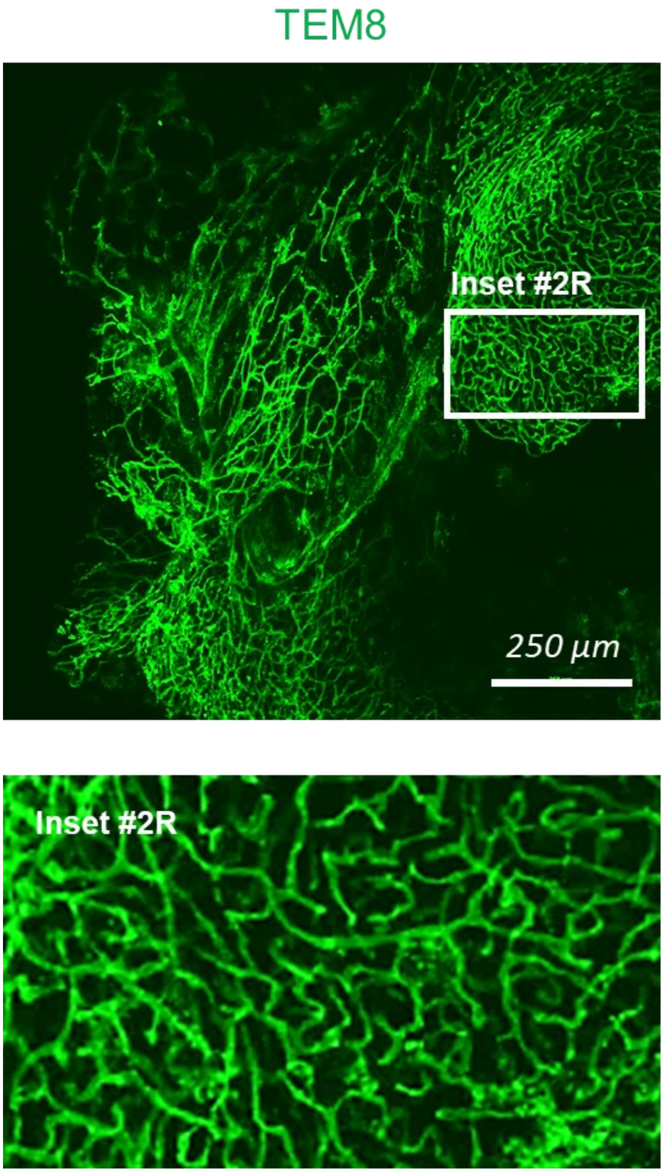


**Figure S12. Confocal Microscopy of a Melanoma Tumor Immunostained for TEM8.** Projection image of a melanoma mass live-immunostained for TEM8 and examined *in toto*. TEM8 is observed along the neovasculature of the tumor organ. The inset (#2R) shows a magnification of the area marked by a box, highlighting the detailed distribution of TEM8.

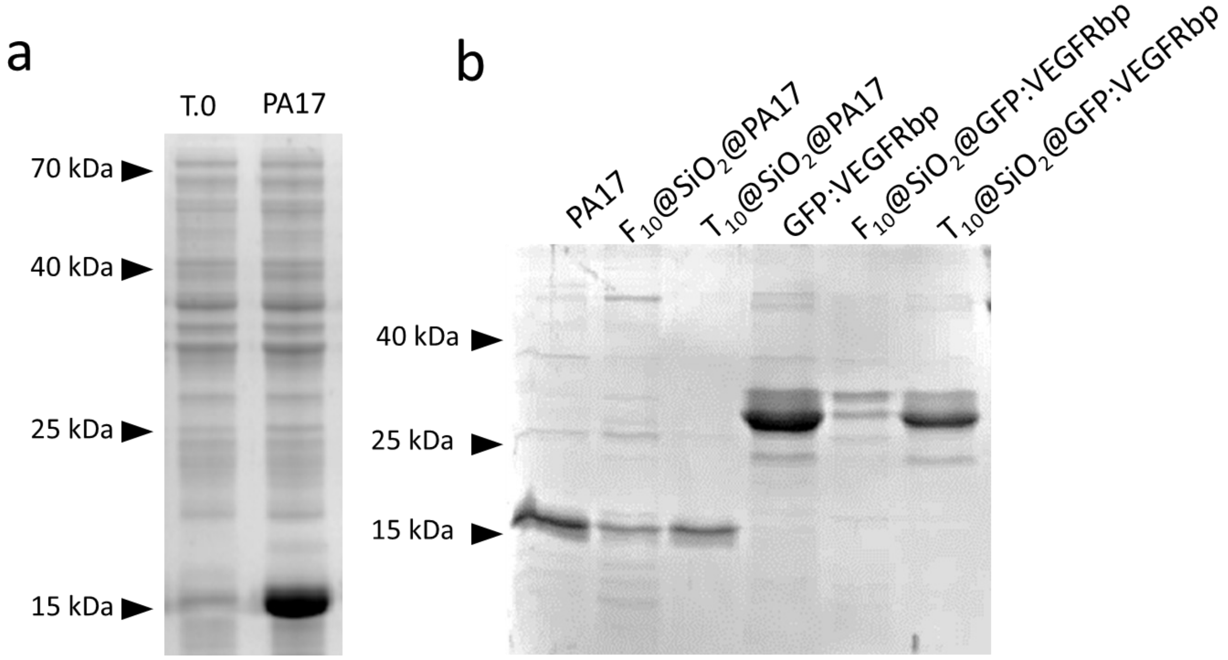


**Figure S13. SDS-PAGE Analysis of the Recombinant Proteins Used. A.** SDS-PAGE gels showing the bacterial extract before and after the overexpression of PA17, demonstrating the presence and abundance of the protein post-expression. **B.** SDS-PAGE gels illustrate the purified ligand proteins and the same proteins after being stripped from the functionalized nanoparticles, as indicated. This demonstrates the successful functionalization of the ligand proteins to the nanoparticles.

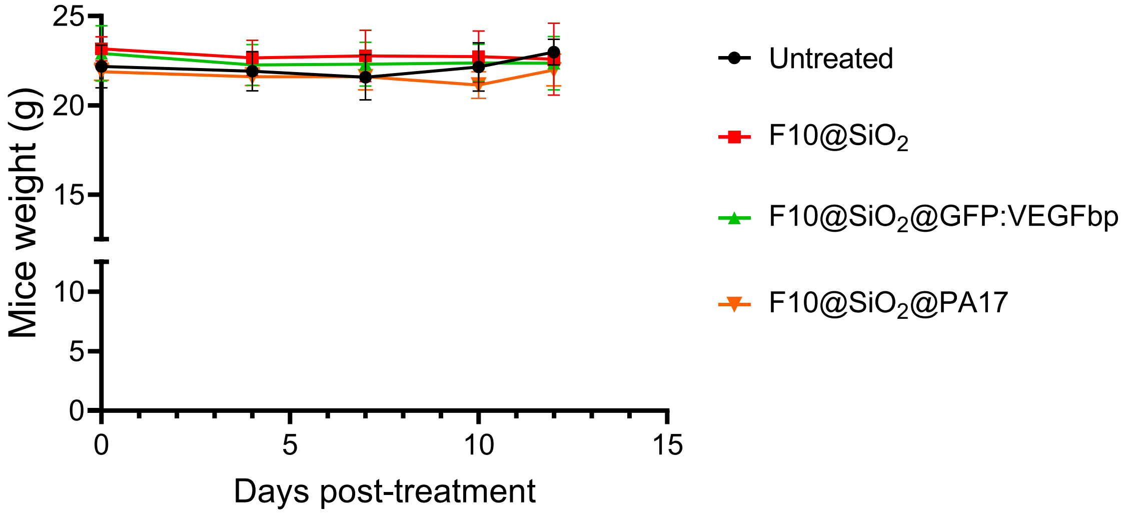


**Figure S14.** Changes in body weight of mice during the period of systemic treatment

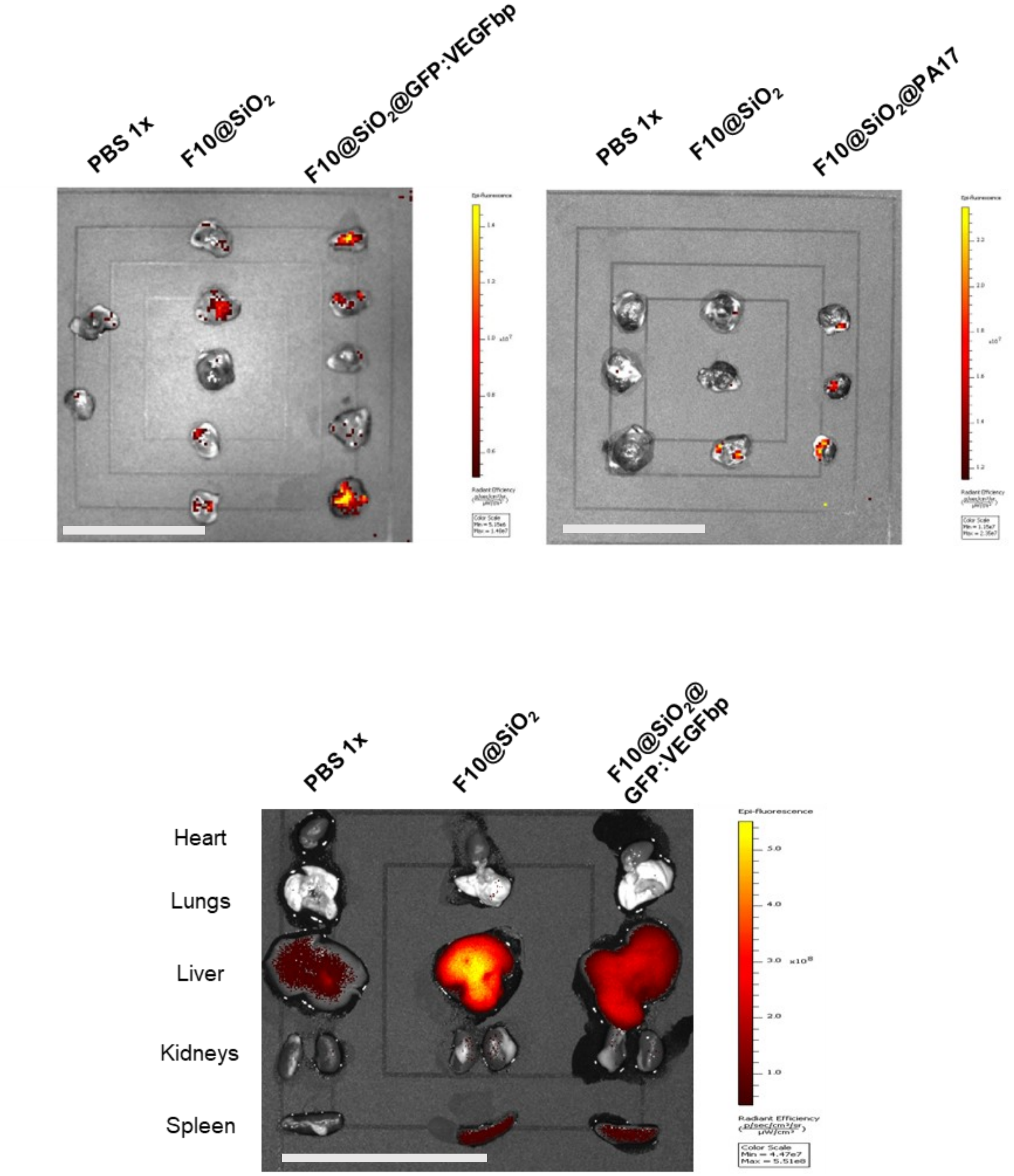


**Figure S15. IVIS® Cy5 Bioimaging of Tumors After Systemic Treatment.** Malignant melanoma tumor organs from mice 72 hours after systemic administration of indicated treatments. The organs were imaged using an IVIS® Biofluorescence imaging system to document the presence of Cy5-labeled prodrug. The bottom image shows the distribution of the fluorescence in other organs of the PBS, and nanoparticle treated mice. Scale bar (grey)=50 mm

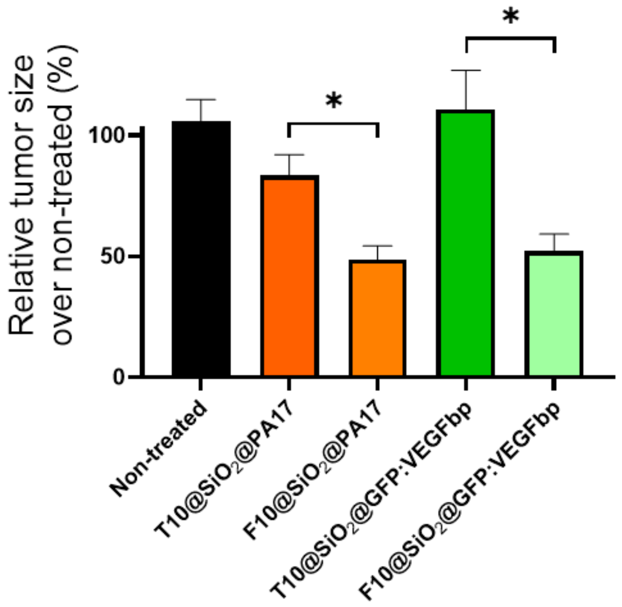


**Figure S16. Antitumoral Effect of the Targeted Nanoparticles.** immobilized ligands GFP:VEGFRbp and PA17 with $T_{10}$@$SiO_2$ and $FdU_{10}$@$SiO_2$ particles. One-way ANOVA with multiple *t*-test comparisons

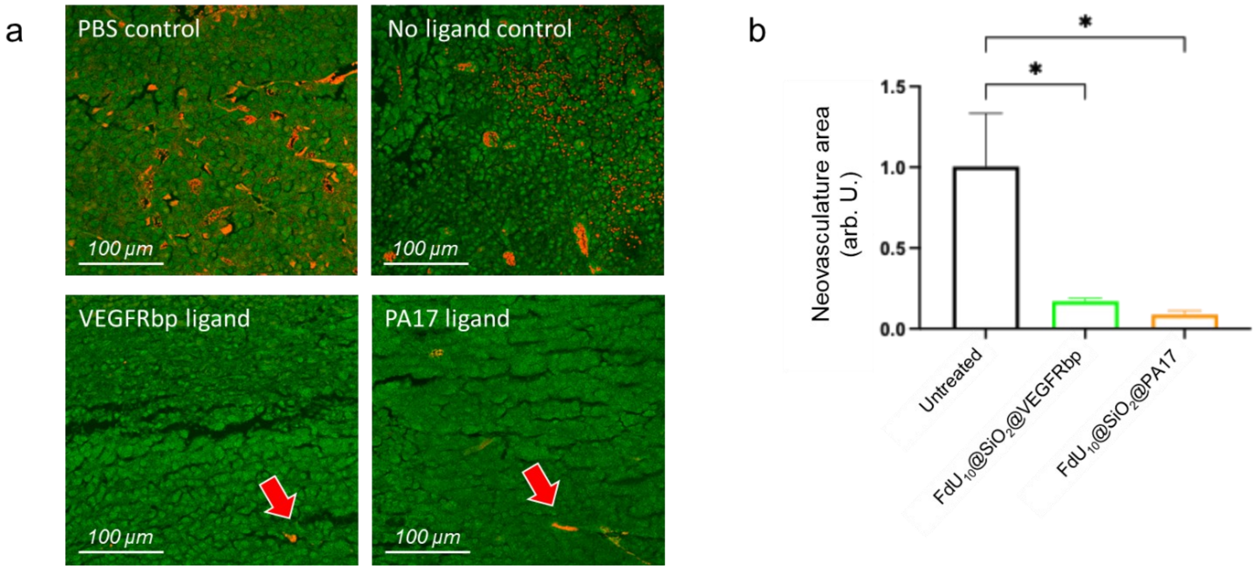


**Figure S17. Intratumoral Neovasculature Analysis Following Systemic Treatments. a.** Confocal microscopy images illustrate tumoral masses embedded in paraffin, sectioned, and stained with hematoxylin-eosin. Blood vessels are identifiable by the presence of erythrocytes, appearing in a distinct bright red color. In treated tumors, only sporadic blood vessels dispersed throughout the tissue are observable, as indicated by arrows. **b**. ImageJ quantification was employed to measure the total surface area occupied by blood vessels in melanoma tumor sections. Comparative analysis revealed a noteworthy reduction in neovasculature in tumors from mice treated with $FdU_{10}$@SiO2@PA17 and $FdU_{10}$@SiO2@VEGFRbp, as opposed to those treated systemically with PBS or non-functionalized nanoparticles. Each group consisted of $n$=3 tumor sections. Statistical analysis using One-way ANOVA with multiple $t$-test comparisons indicated significance (*$p$<0.05).
